# A Master Autoantigen-ome Links Alternative Splicing, Female Predilection, and COVID-19 to Autoimmune Diseases

**DOI:** 10.1101/2021.07.30.454526

**Authors:** Julia Y. Wang, Michael W. Roehrl, Victor B. Roehrl, Michael H. Roehrl

**Affiliations:** Curandis, New York, USA; Department of Pathology, Memorial Sloan Kettering Cancer Center, New York, USA

## Abstract

Chronic and debilitating autoimmune sequelae pose a grave concern for the post-COVID-19 pandemic era. Based on our discovery that the glycosaminoglycan dermatan sulfate (DS) displays peculiar affinity to apoptotic cells and autoantigens (autoAgs) and that DS-autoAg complexes cooperatively stimulate autoreactive B1 cell responses, we compiled a database of 751 candidate autoAgs from six human cell types. At least 657 of these have been found to be affected by SARS-CoV-2 infection based on currently available multi-omic COVID data, and at least 400 are confirmed targets of autoantibodies in a wide array of autoimmune diseases and cancer. The autoantigen-ome is significantly associated with various processes in viral infections, such as translation, protein processing, and vesicle transport. Interestingly, the coding genes of autoAgs predominantly contain multiple exons with many possible alternative splicing variants, short transcripts, and short UTR lengths. These observations and the finding that numerous autoAgs involved in RNA-splicing showed altered expression in viral infections suggest that viruses exploit alternative splicing to reprogram host cell machinery to ensure viral replication and survival. While each cell type gives rise to a unique pool of autoAgs, 39 common autoAgs associated with cell stress and apoptosis were identified from all six cell types, with several being known markers of systemic autoimmune diseases. In particular, the common autoAg UBA1 that catalyzes the first step in ubiquitination is encoded by an X-chromosome escape gene. Given its essential function in apoptotic cell clearance and that X-inactivation escape tends to increase with aging, UBA1 dysfunction can therefore predispose aging women to autoimmune disorders. In summary, we propose a model of how viral infections lead to extensive molecular alterations and host cell death, autoimmune responses facilitated by autoAg-DS complexes, and ultimately autoimmune diseases. Overall, this master autoantigen-ome provides a molecular guide for investigating the myriad of autoimmune sequalae to COVID-19 and clues to the rare but reported adverse effects of the currently available COVID vaccines.

## Introduction

Autoimmune disorders are an important feature of the disease manifestations of COVID-19 and long-COVID syndromes. Based on the insights we gained from numerous COVID-related autoantigens (autoAgs) and their associated cellular process and pathways [1–5], we propose a model to explain how viral infections in general and SARS-CoV-2 in particular can lead to a wide array of autoimmune diseases (Figure 1). We illustrate how viral infections lead to extensive molecular alterations in the host cell, host cell death and tissue injury, autoimmune reactions, and the eventual development of autoimmune diseases.

**Fig. 1.**
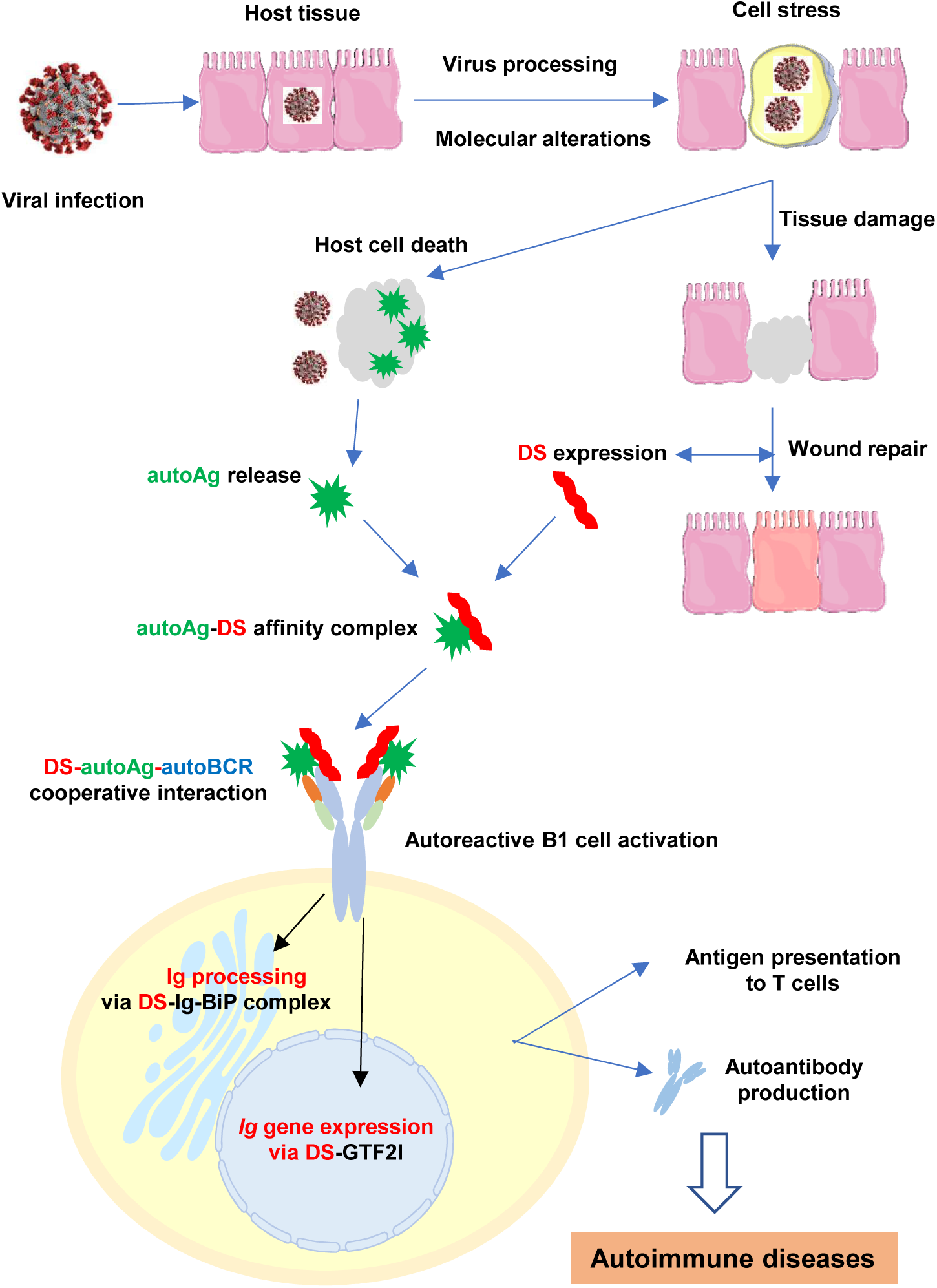
A model on how viral infections lead to autoimmune diseases. Viral infections induce extensive host molecular changes, cell death, and tissue damage. AutoAgs shed from apototic cells form affinity complexes with DS that is overexpressed in the wound area. Cooperative binding of DS-autoAg complexes to autoBCRs activate autoreactive B1 cells. Once internalized via autoBCR, DS engages Ig-processing complexes in the ER and GTF2I in the nucleus to facilitate Ig production. Activated B1 cells secrete autoantibodies and may also present autoAgs to autoreactive T cells, which then leads to autoimmune diseases.

During infections, opportunistic viruses have to hijack the host cell machinery in order to transcribe and translate the viral genes, synthesize viral proteins with correct polypeptide folding and post-translational modifications, and assemble viral particles. At the same time, viruses have to manipulate the host’s immune defense to avoid elimination. This intricate host-virus symbiosis is accomplished by extensive alterations of host molecules and reprogramming of host molecular networks. The infected host cells undergo extreme stress and ultimately die, which releases altered molecules (i.e., potential autoAgs) that the immune system may recognize as non-self. In response, the host also synthesizes a cascade of molecules such as dermatan sulfate (DS) to facilitate wound healing and dead cell clearance.

We have discovered previously that DS possesses peculiar affinity for apoptotic cells and their released autoAgs [6–9]. DS, a major component of the extracellular matrix and connective tissue, is increasingly expressed during tissue injury and accumulates in wound areas [1, 10]. Because of their affinity, DS and autoAgs form macromolecular complexes which cooperatively activate autoreactive B1 cells. AutoAg-DS complexes may activate B1 cells via a dual binding mode, i.e., with autoAg binding to the variable region of the B1 cell’s autoBCR and DS binding to the heavy chain of the autoBCR. Upon entering B1 cells, DS may regulate immunoglobulin (Ig) production by engaging the Ig-processing complex in the endoplasmic reticulum and the transcription factor GTF2I necessary for Ig gene expression [8, 9]. AutoAg-DS affinity therefore defines a unifying biochemical and immunological property of autoAgs: any self-molecule possessing DS-affinity has a high propensity to become autoantigenic, and this has led to the identification of numerous autoAgs [7, 11–13].

To gain a better understanding of autoimmune sequelae due to COVID-19, we present a master autoantigen atlas of over 750 potential autoAgs identified from six human cell types [1, 2, 4, 5, 7, 11]. These autoAgs show significant correlation with pathways and processes that are crucial in viral infection and mRNA vaccine action, reveal common autoAgs associated with apoptosis and cell stress which may serve as markers for systemic autoimmune diseases, and provide a detailed molecular map for understanding and for investigating diverse autoimmune sequalae of COVID-19 and potential rare side-effects to viral vector- and mRNA-based vaccines. For the first time, we reveal intriguing features of autoAgs and their coding genes. Furthermore, we discuss how UBA1 (or UBE1, ubiquitin-like modifier-activating enzyme 1), an autoAg found overexpressed in SARS-CoV-2 infection, may predispose aging females to autoimmune disorders.

## Results and Discussion

### The master autoantigen-ome

To understand the diversity of autoimmune diseases, we were curious to know how many autoAgs possibly exist. A total of 751 potential autoAgs were identified (Table 1) when we combined all DS-affinity autoAgs profiled from six human cell lines, namely, HFL1 fetal lung fibroblasts, HEp2 fibroblasts, A549 lung epithelial cells, HS-Sultan and Wil2-NS B-lymphoblasts, and Jurkat T-lymphoblasts. Extensive literature searches confirmed that at least 400 of these proteins (53.3%) have been reported as targets of autoantibodies in a wide variety of autoimmune diseases and cancer (see autoAg confirmation references in Table 1). The majority of unconfirmed or putative autoAgs are isoforms of or structurally similar to reported autoAgs and are yet-to-confirmed autoAgs. For example, 56 ribosomal proteins were identified by DS-affinity, but only 22 are thus far confirmed autoAgs; but given their structural similarity and shared epitopes, it is likely that most if not all of the 56 ribosomal proteins are likely true autoAgs awaiting further confirmation.

**Table 1.**
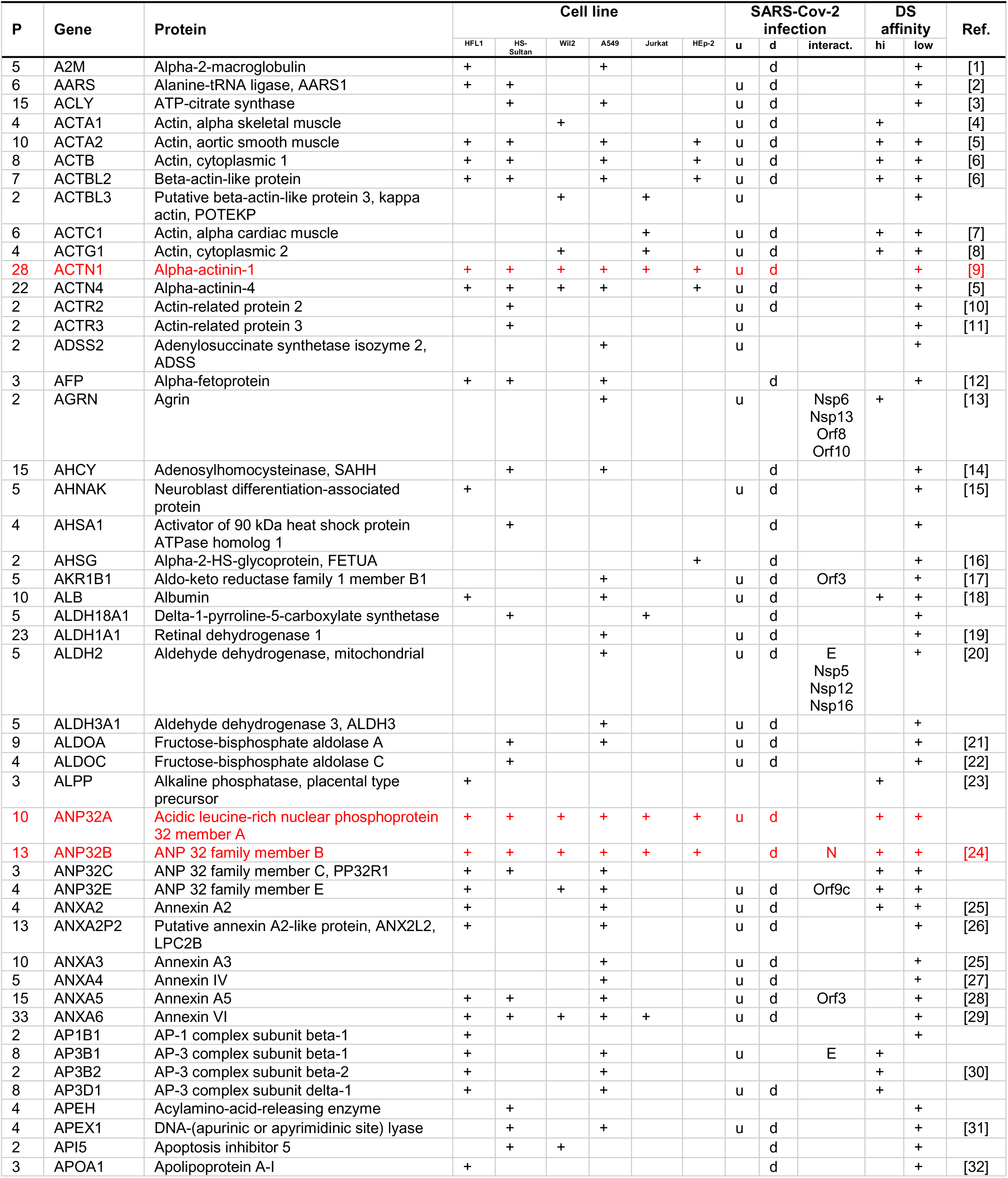
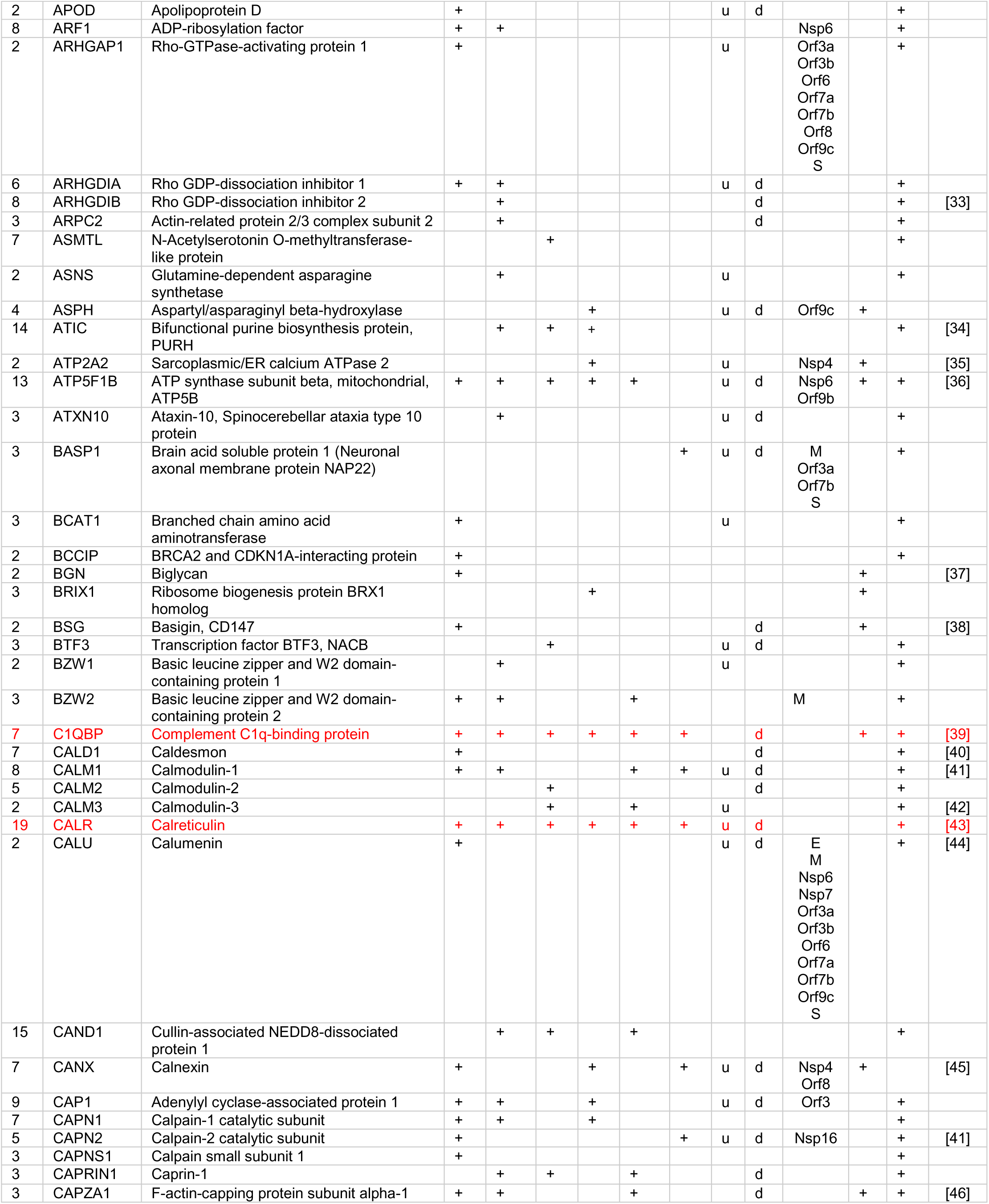
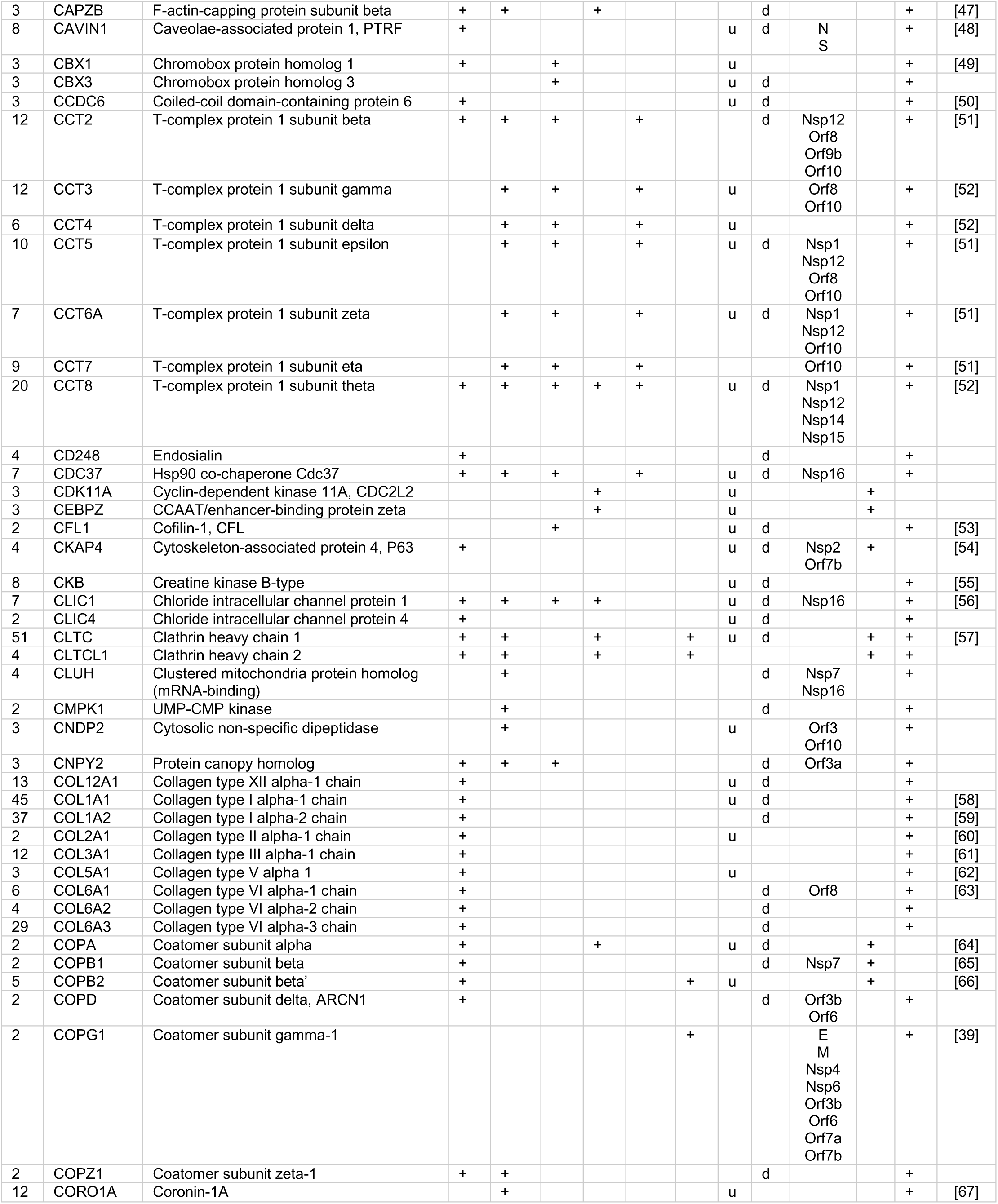
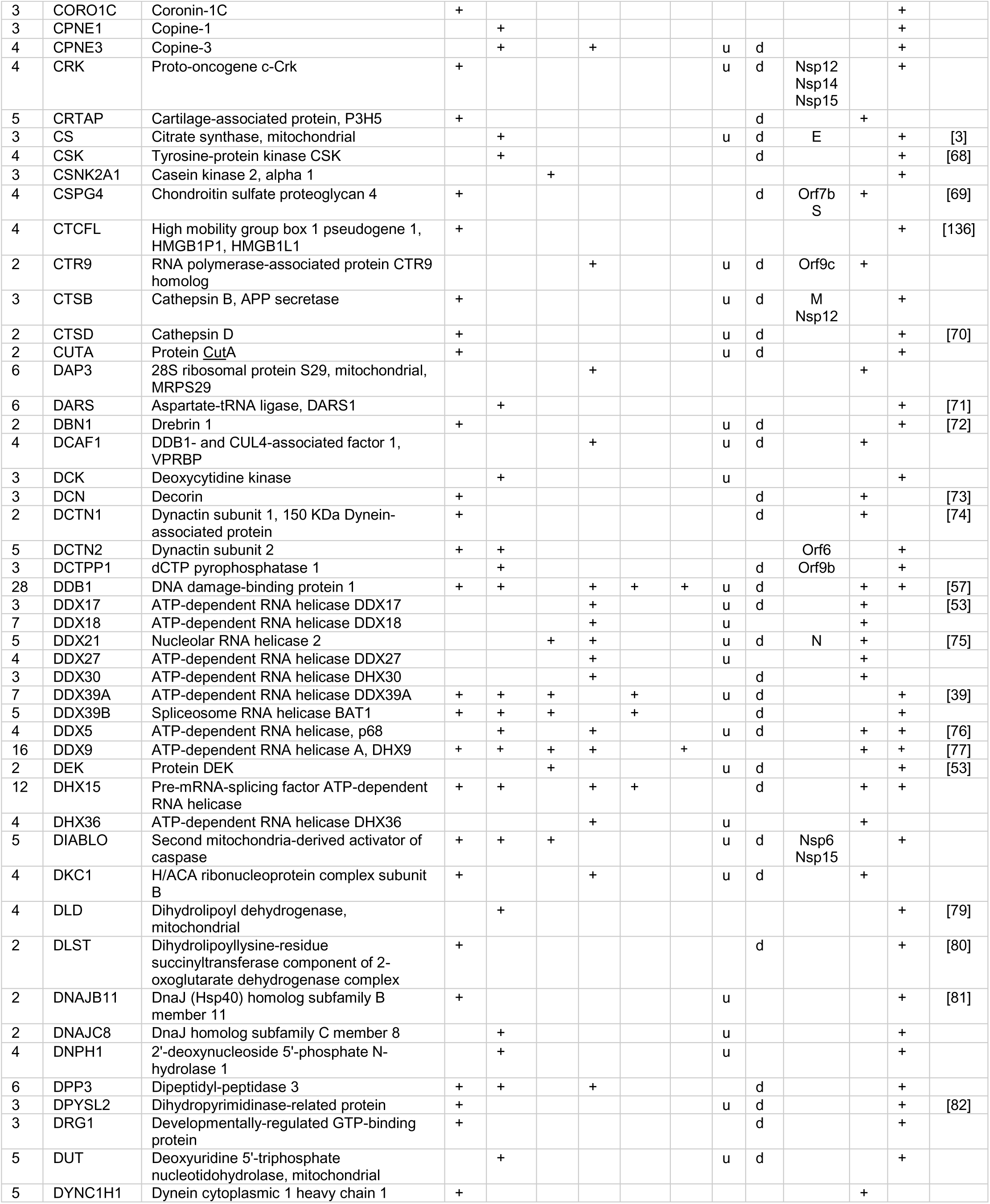
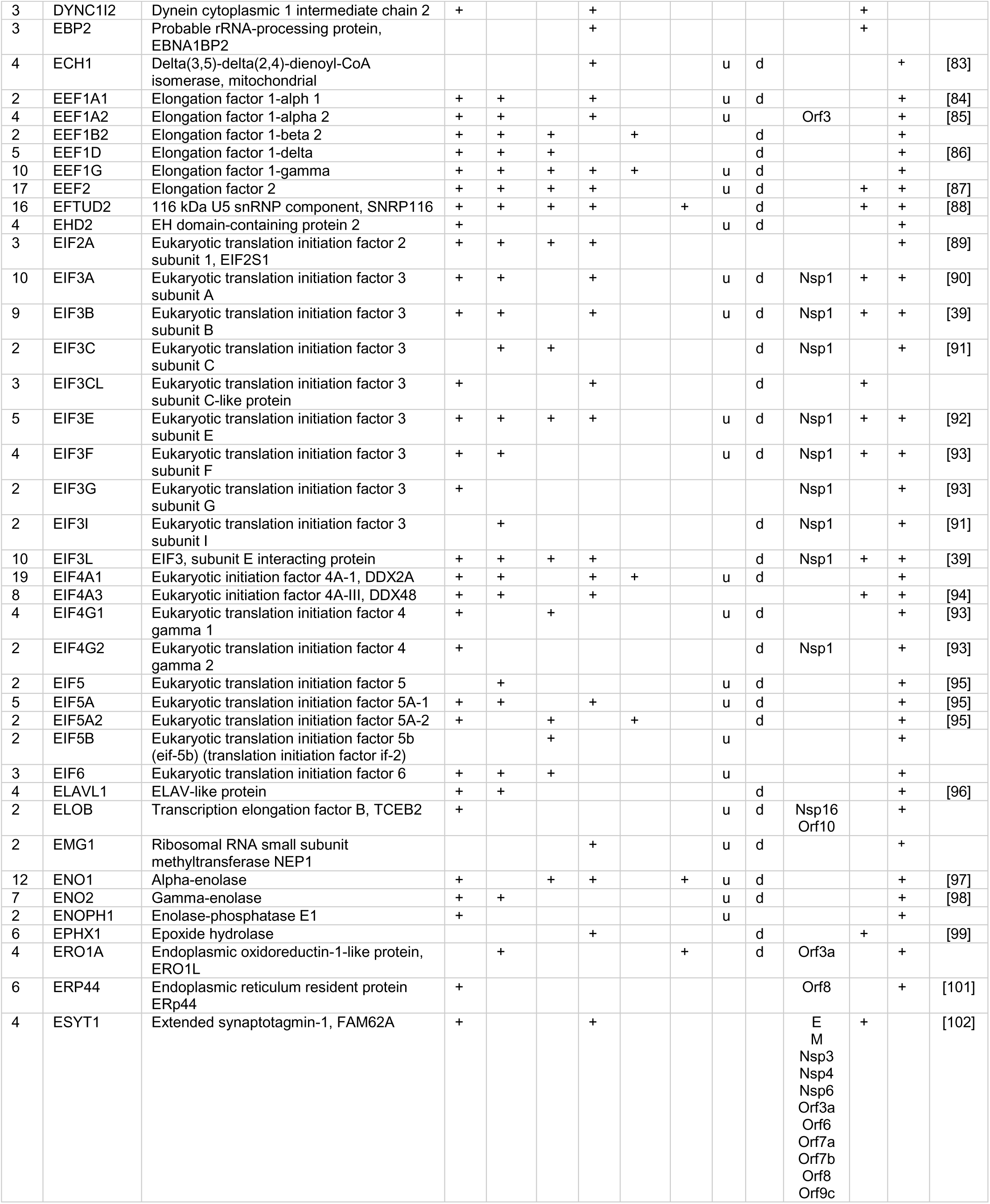
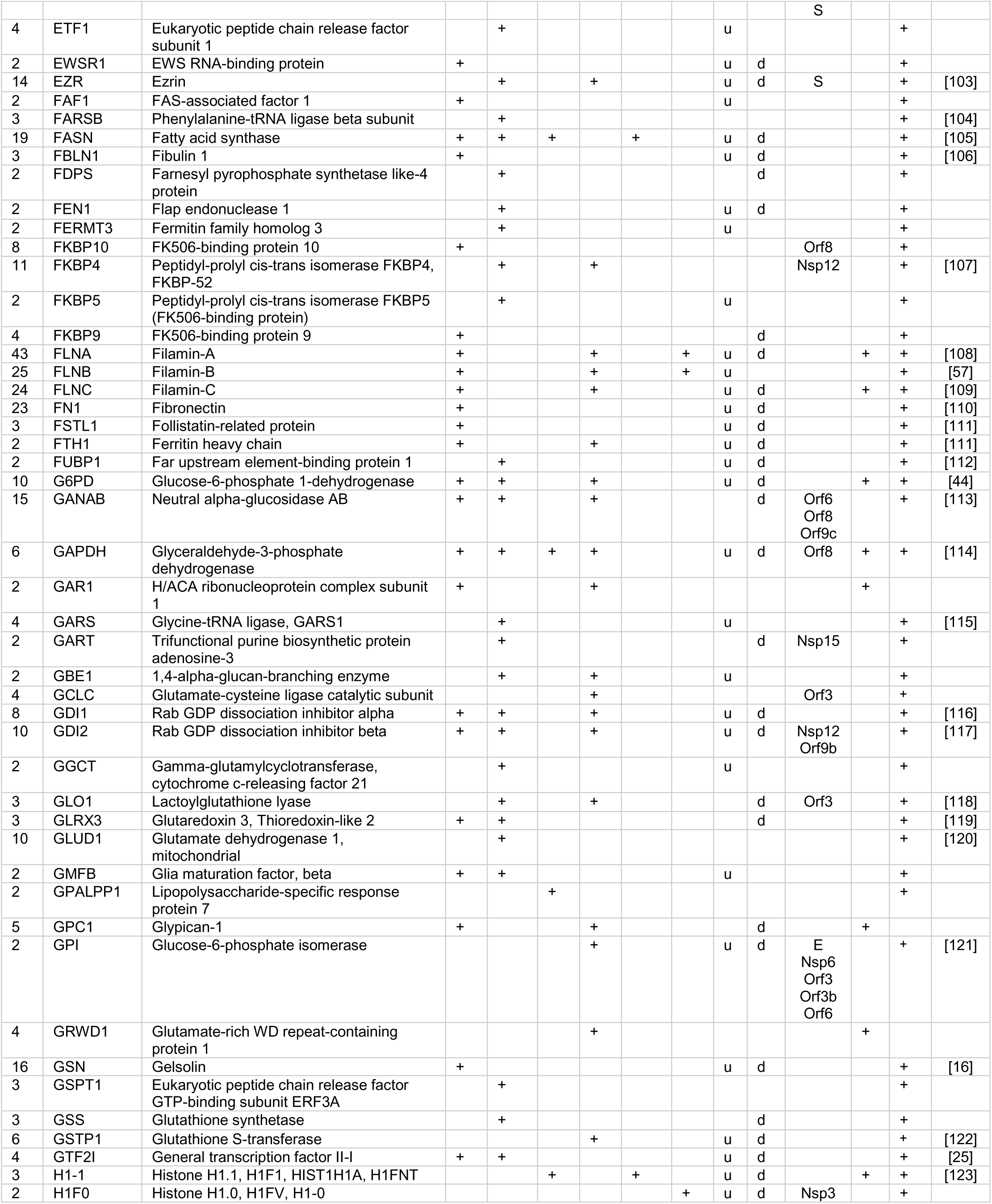
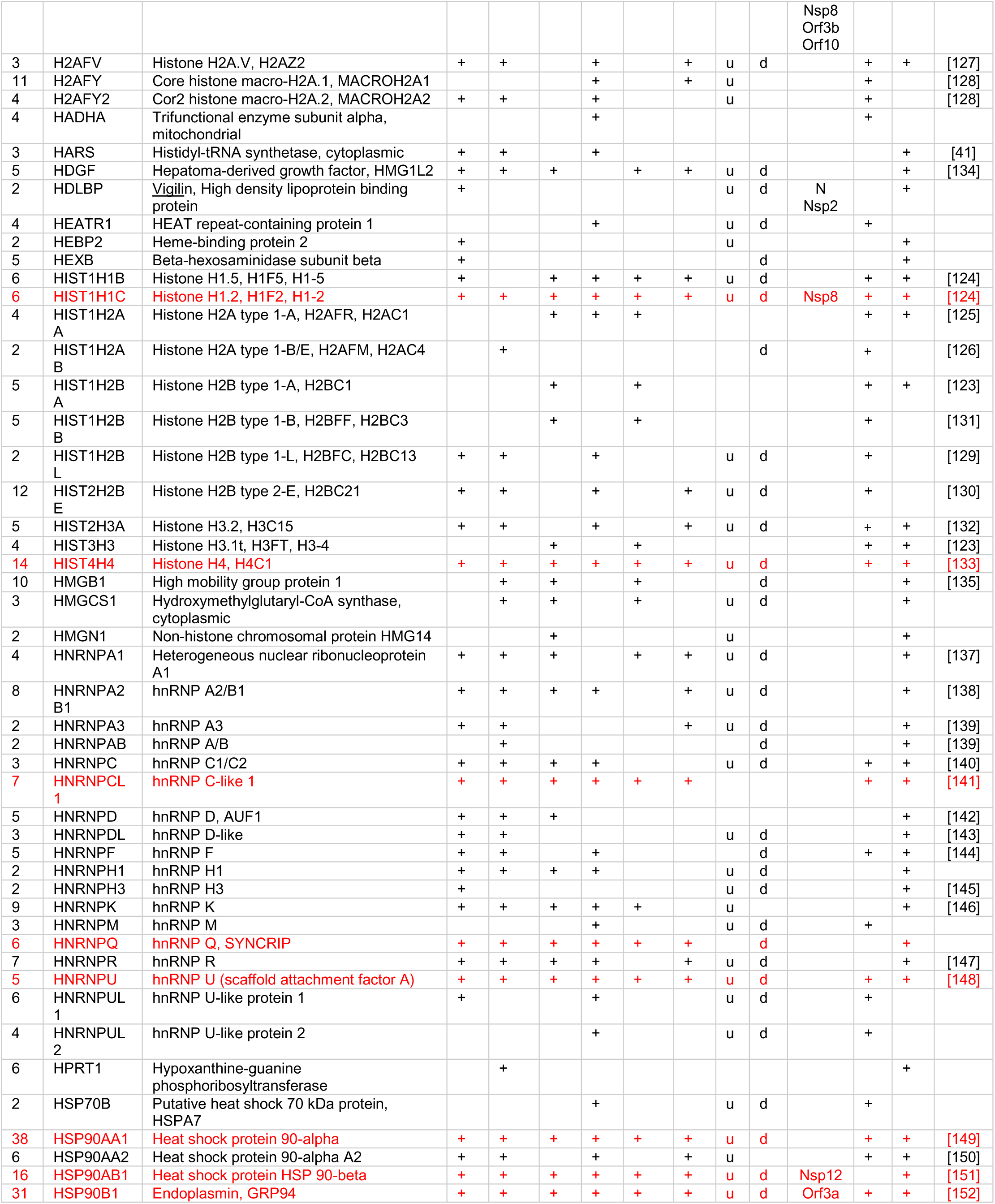
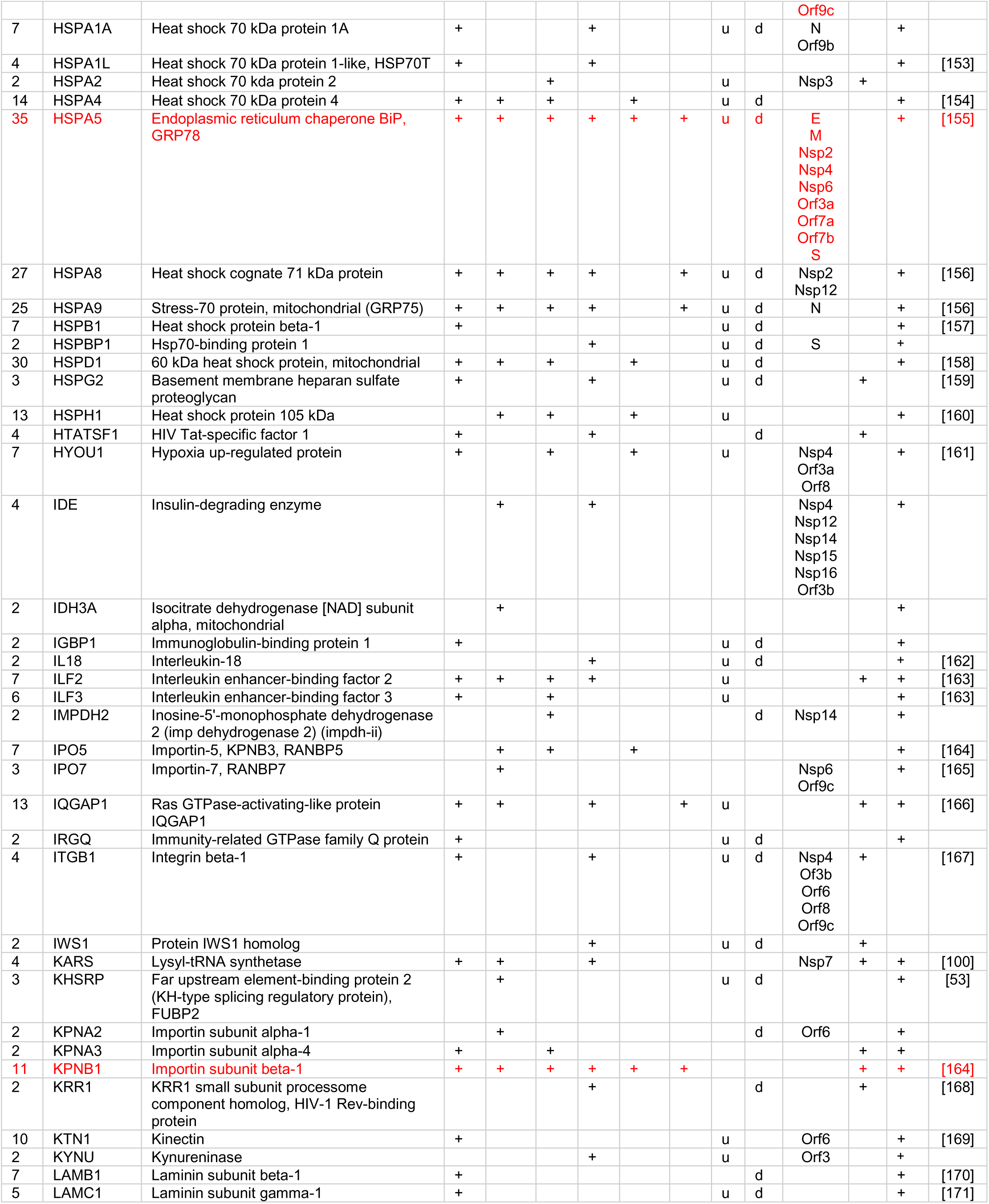
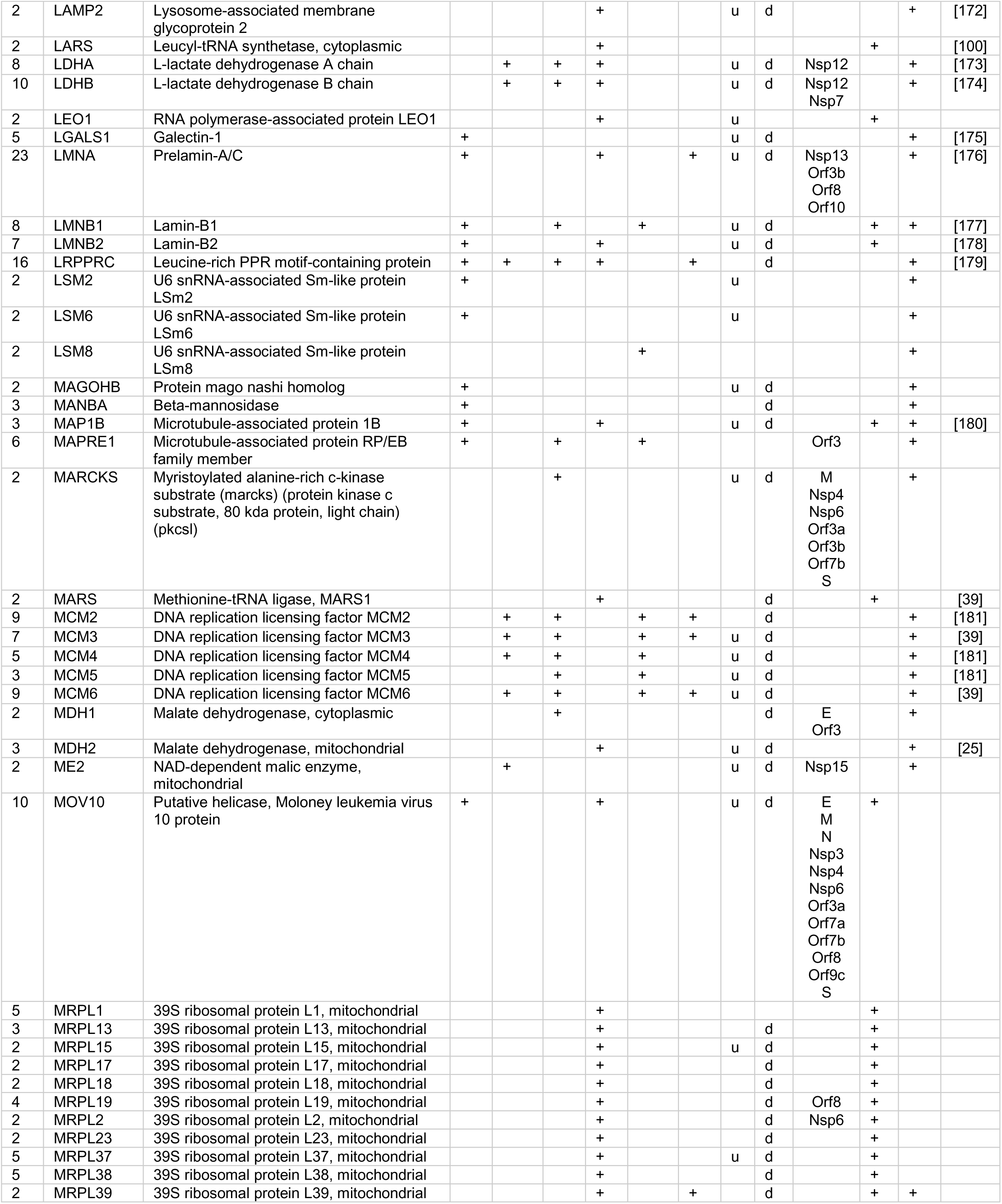
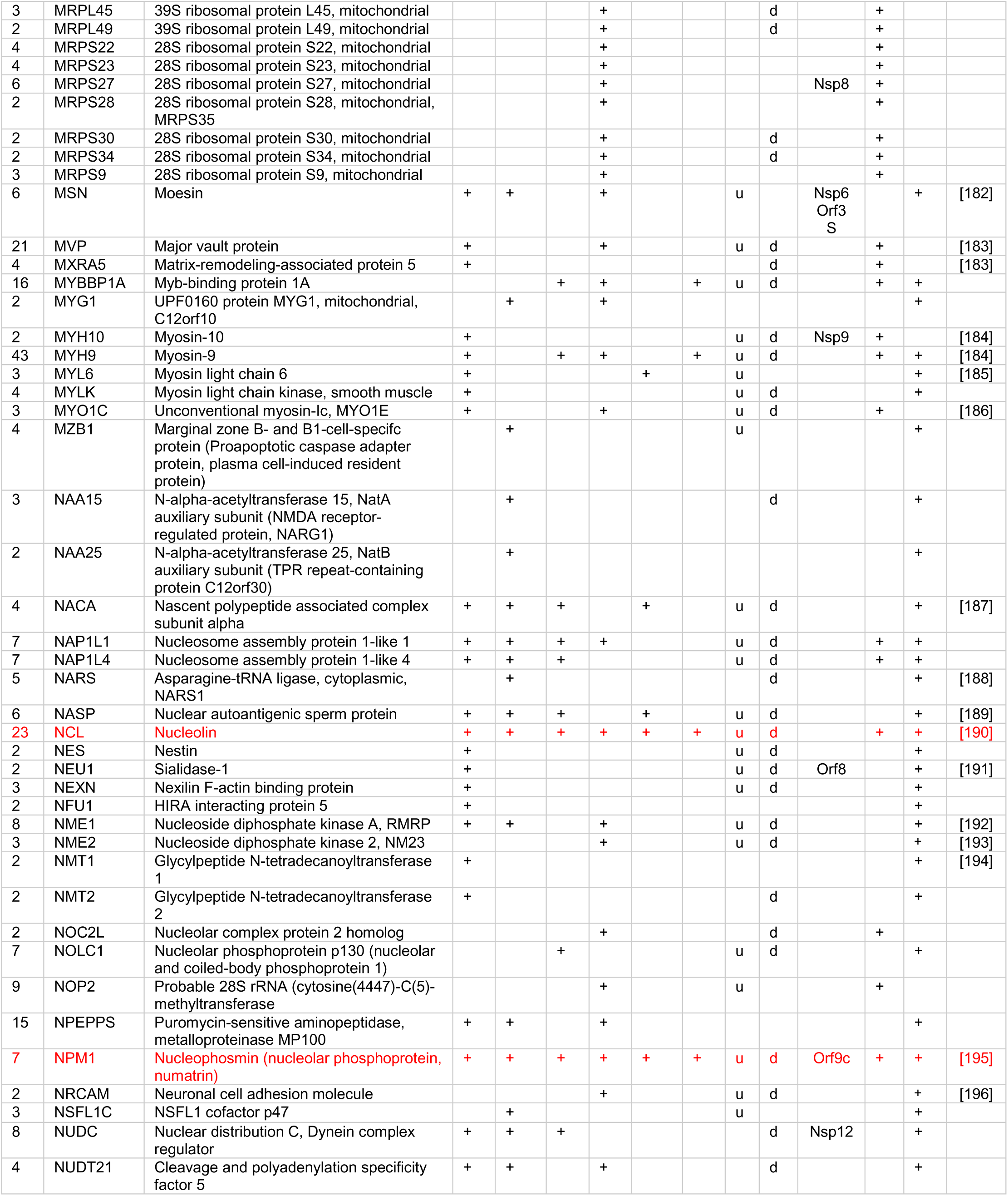
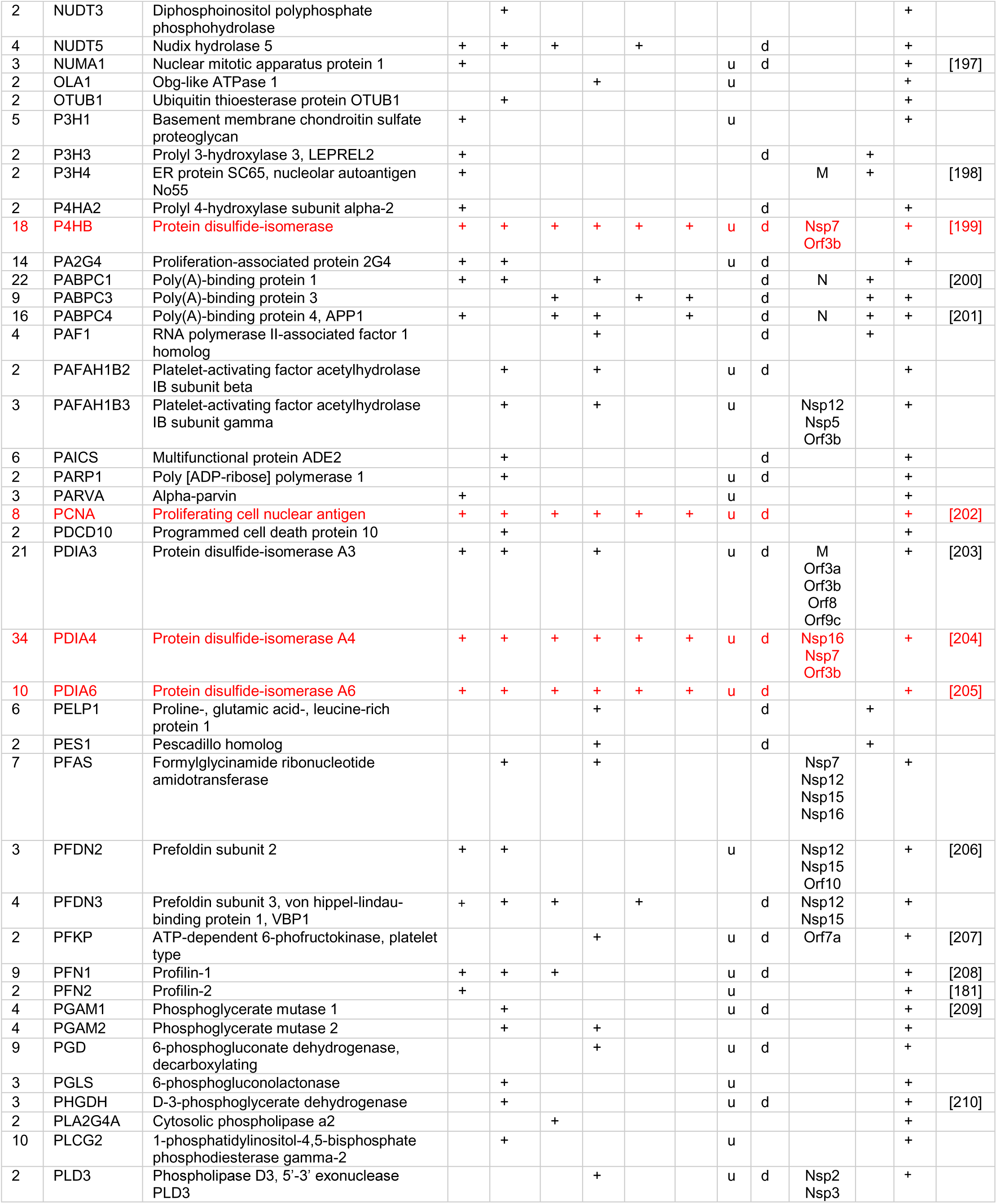
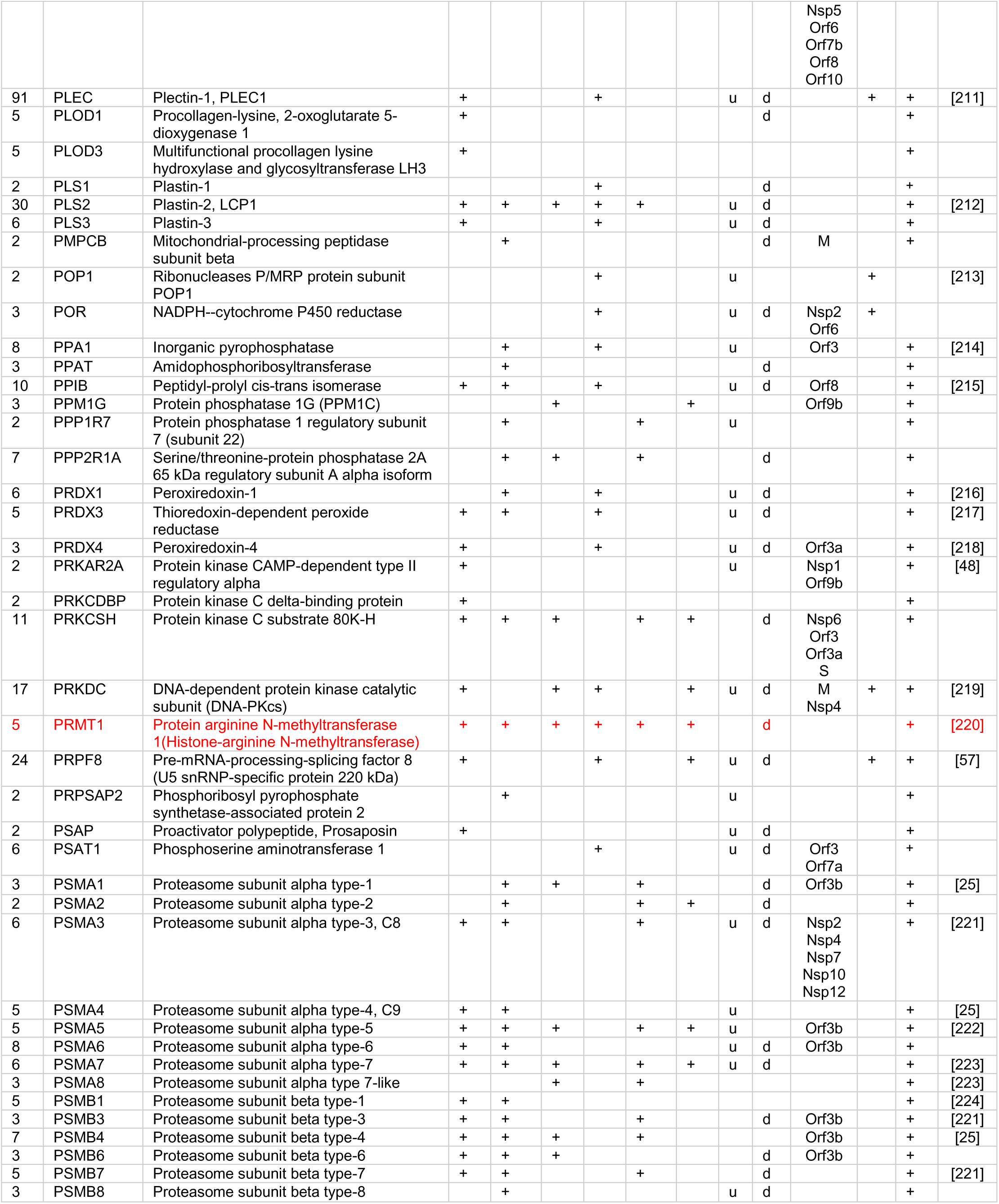
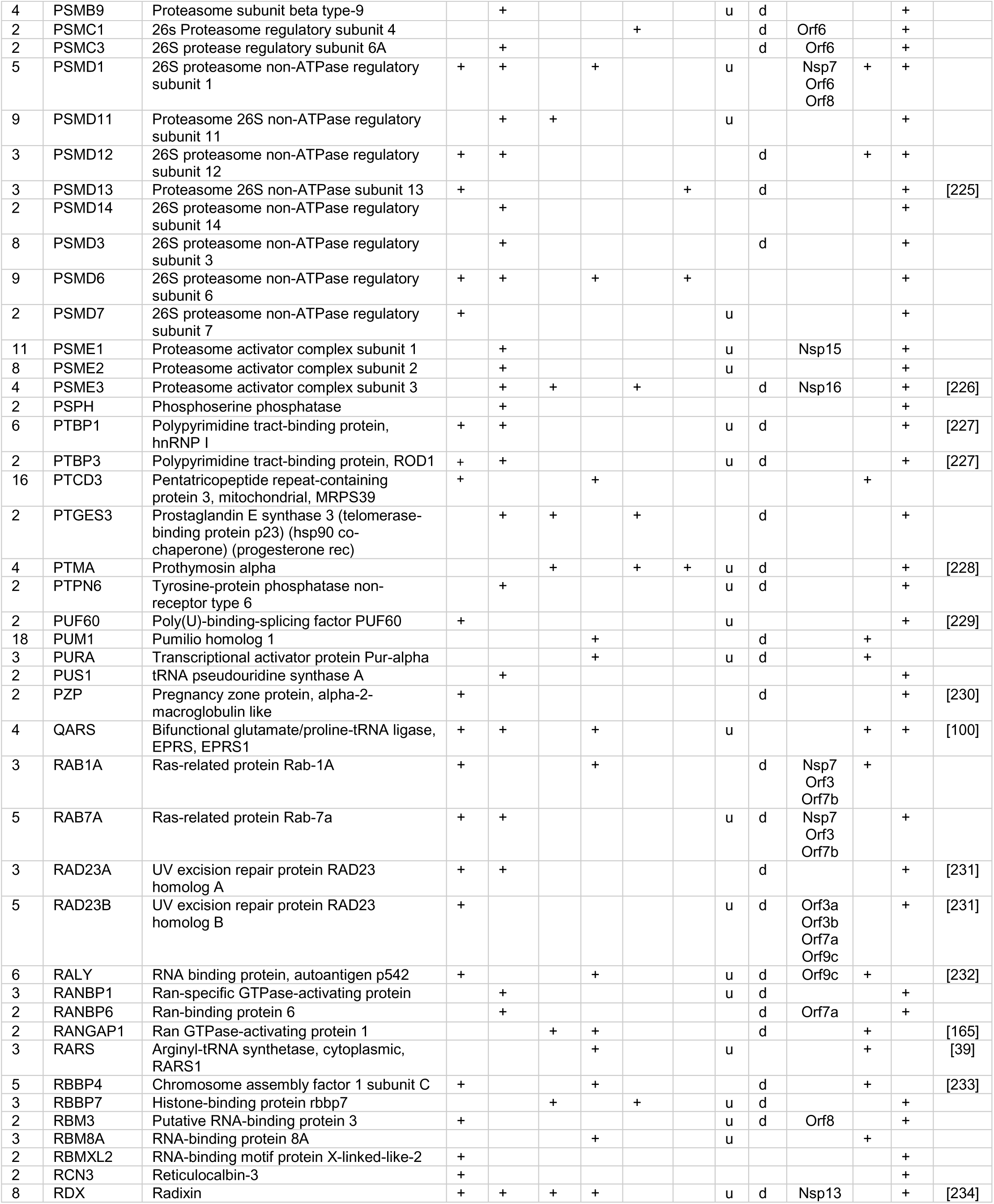
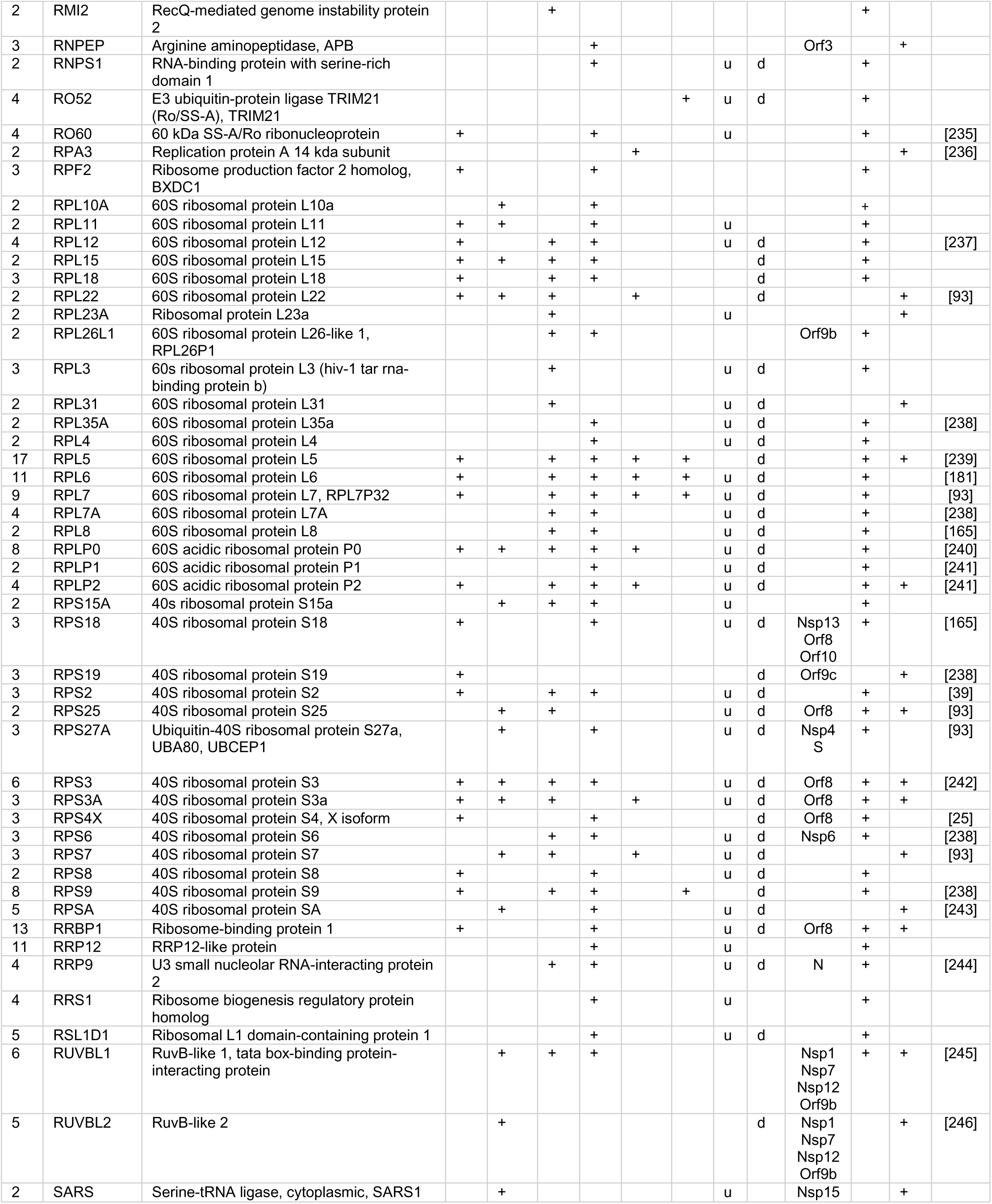
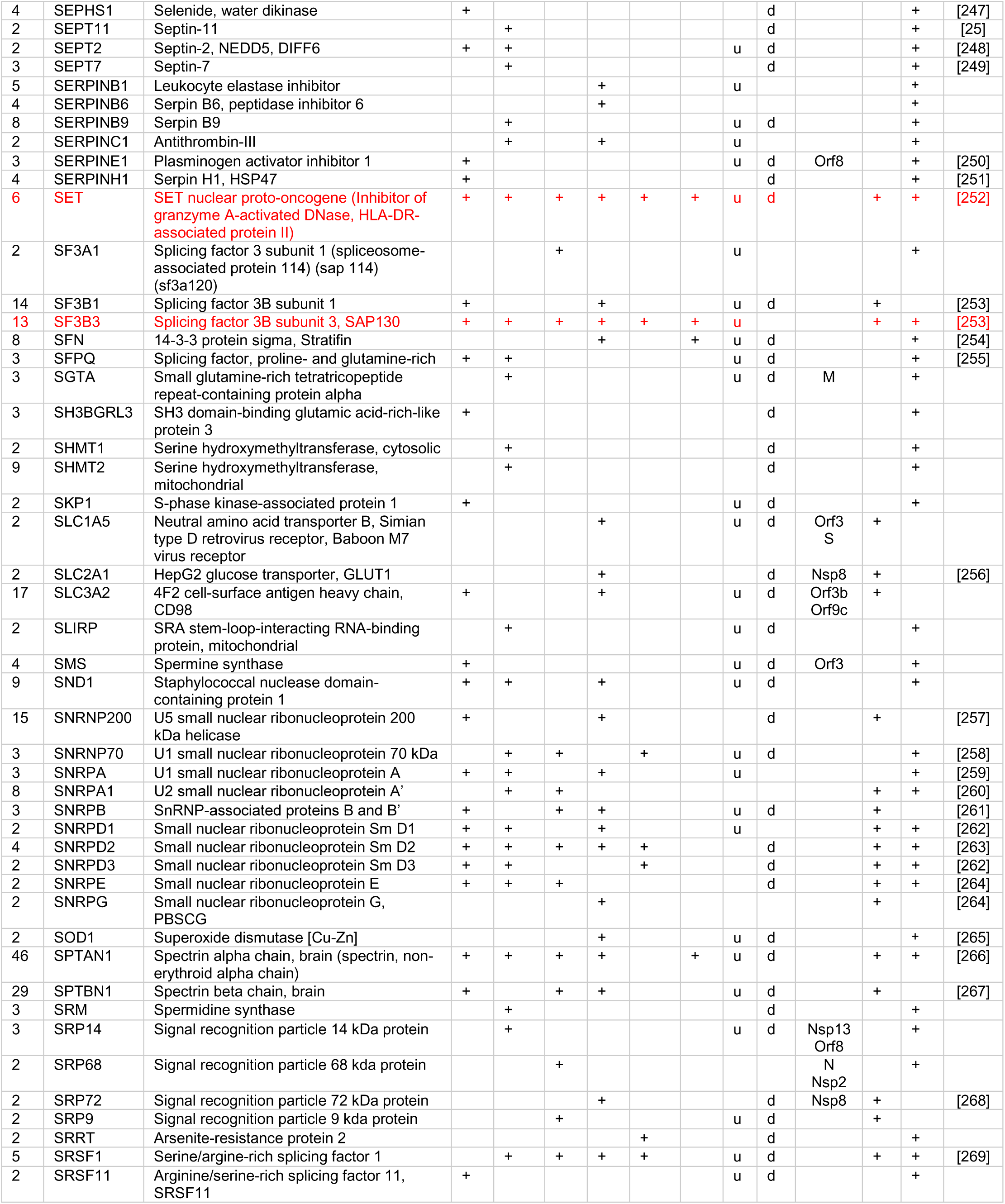
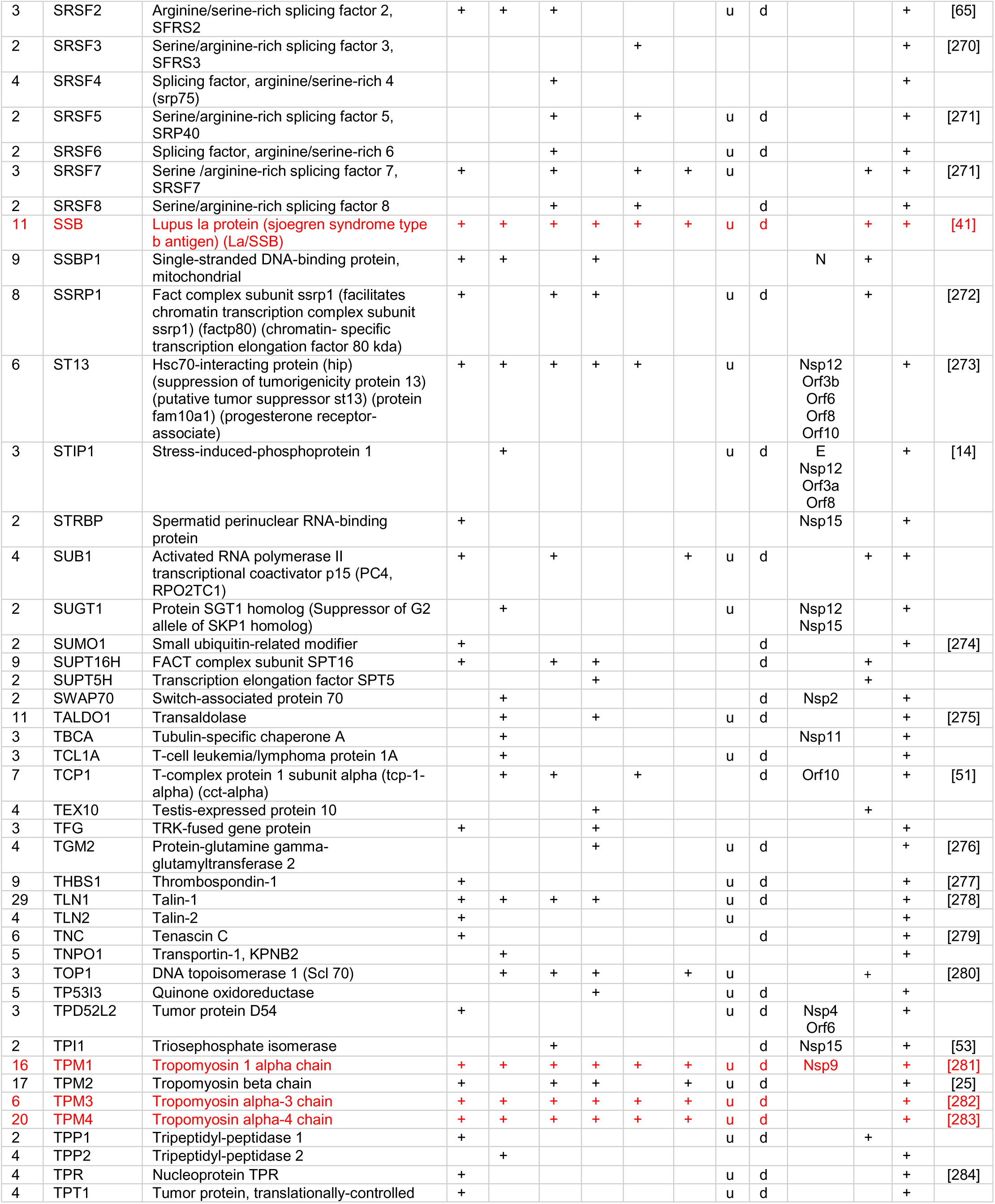
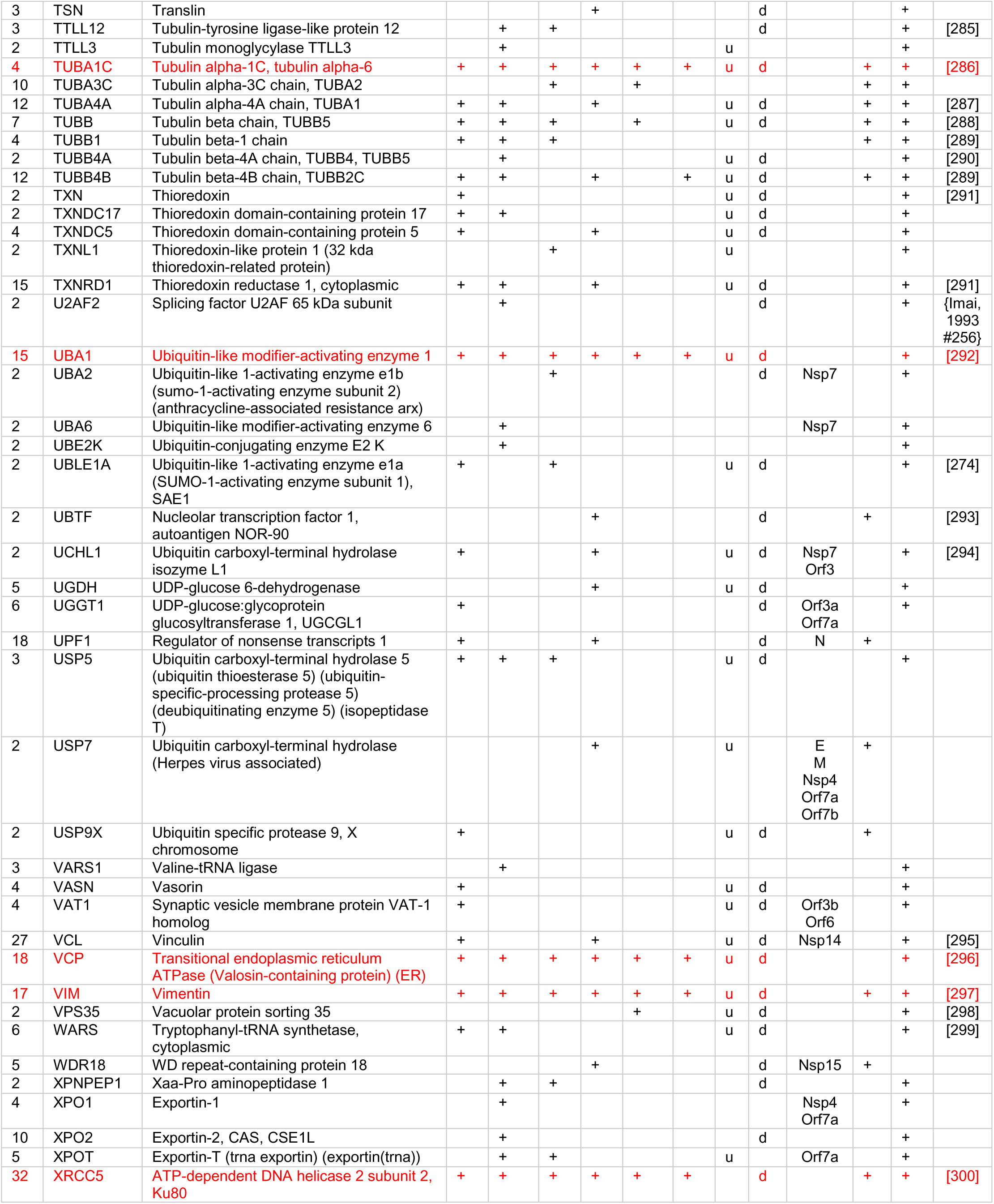
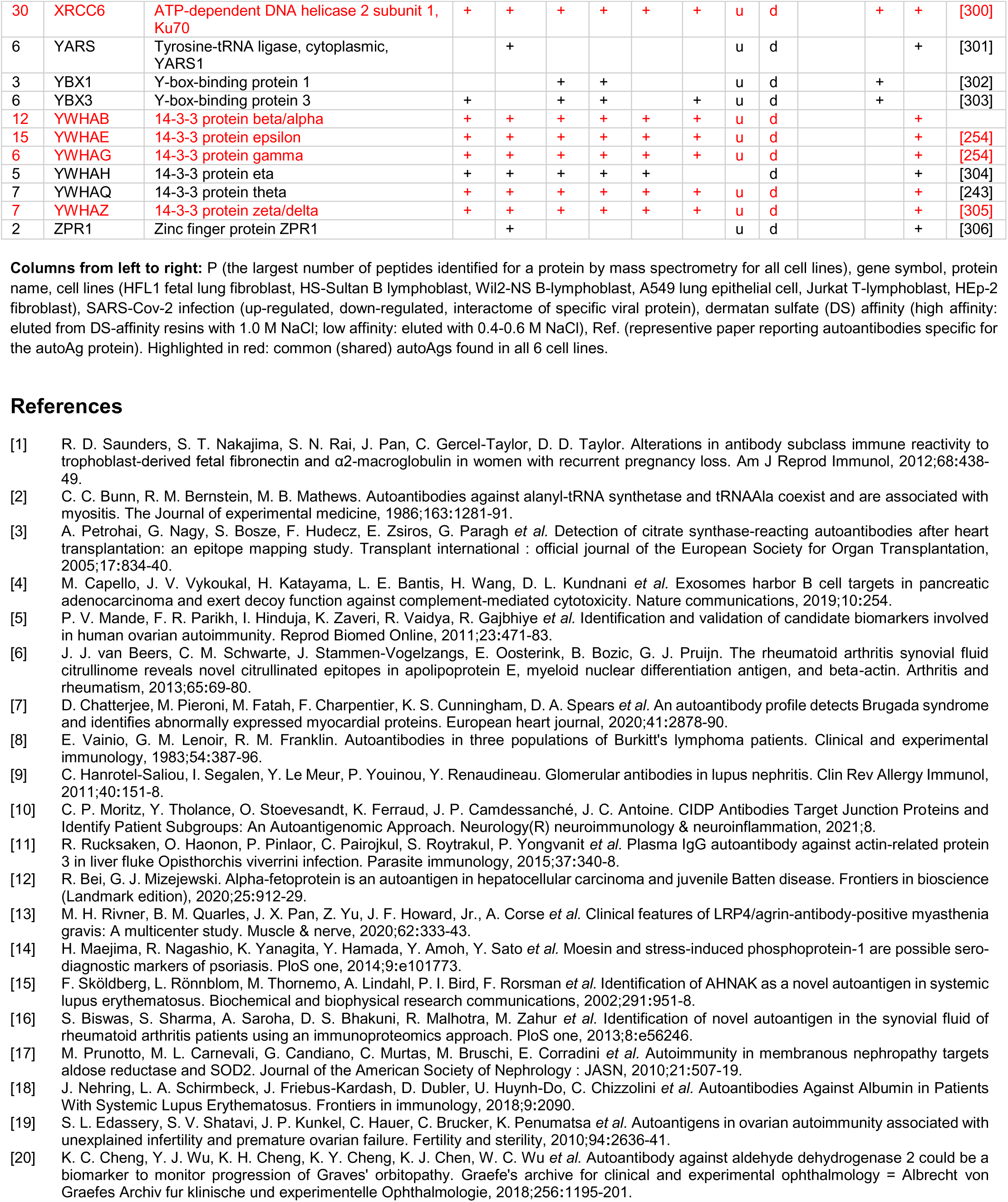
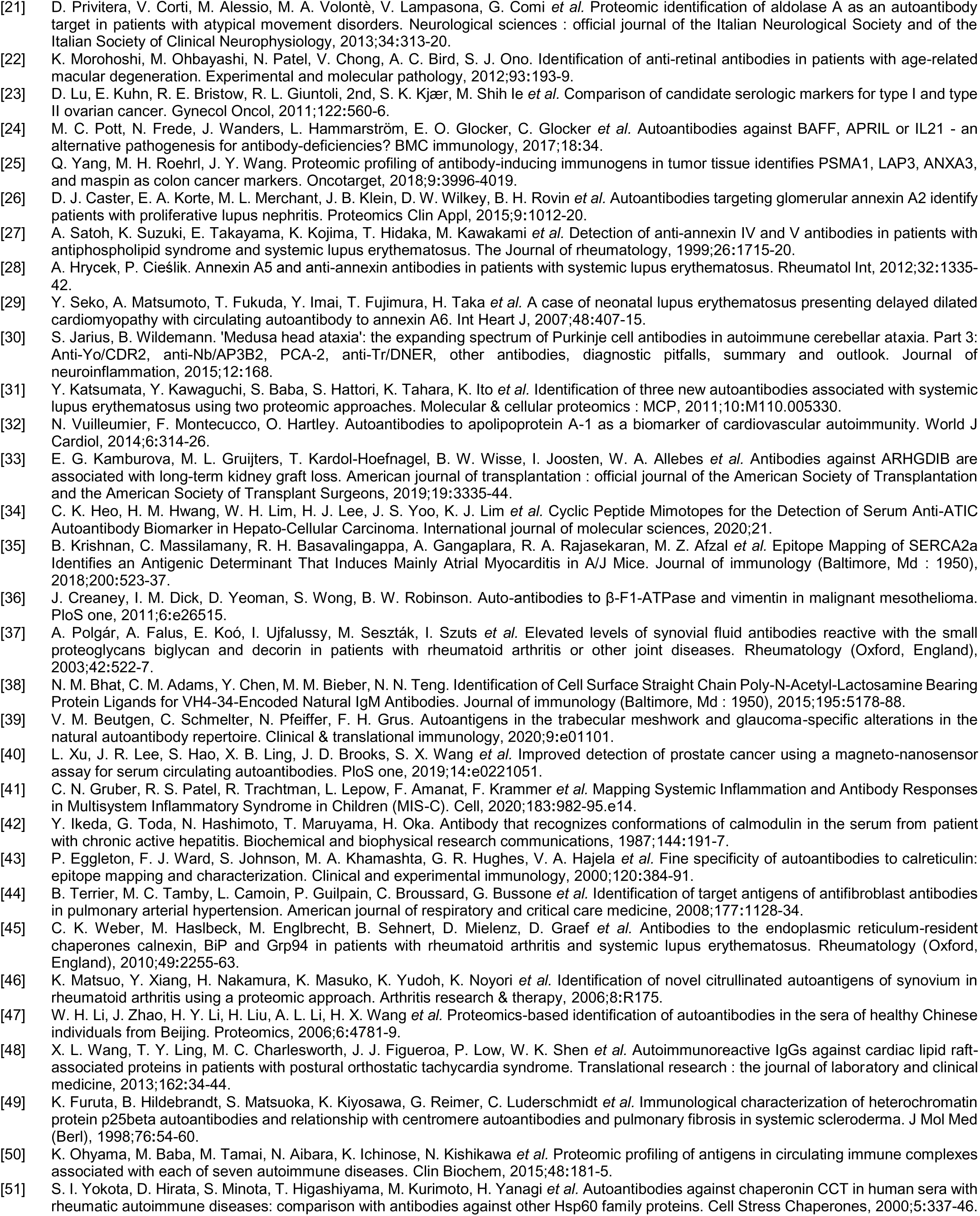
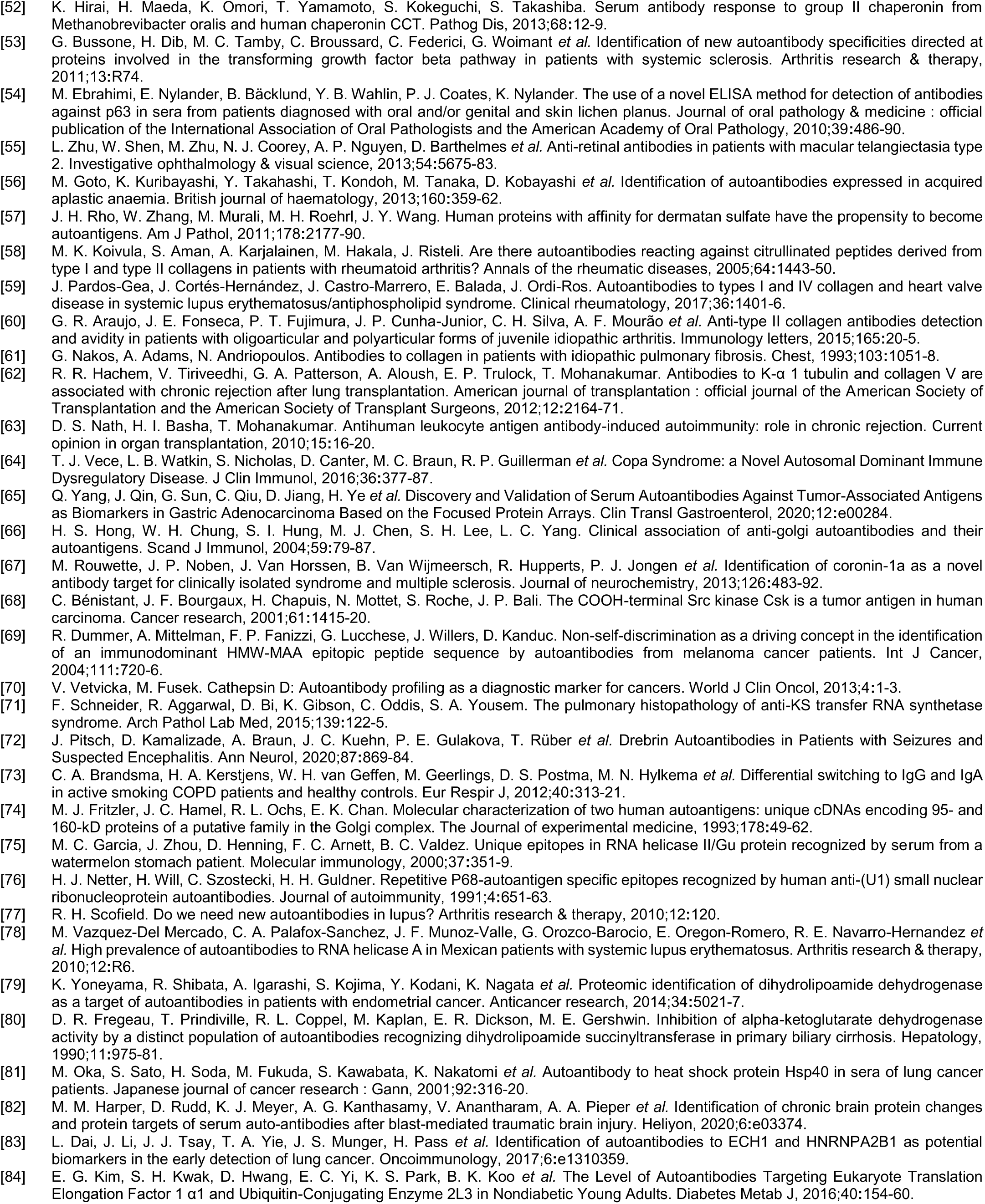
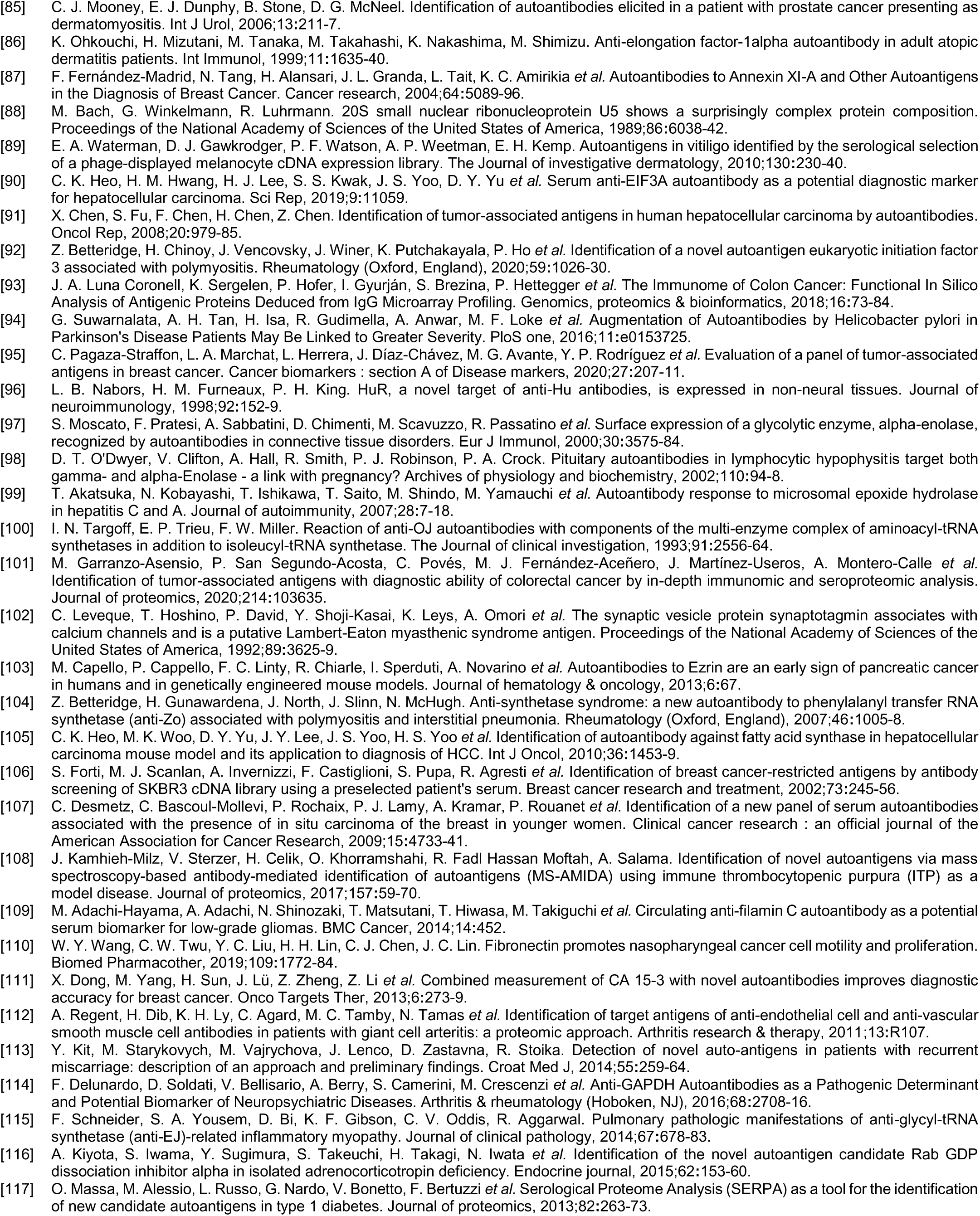
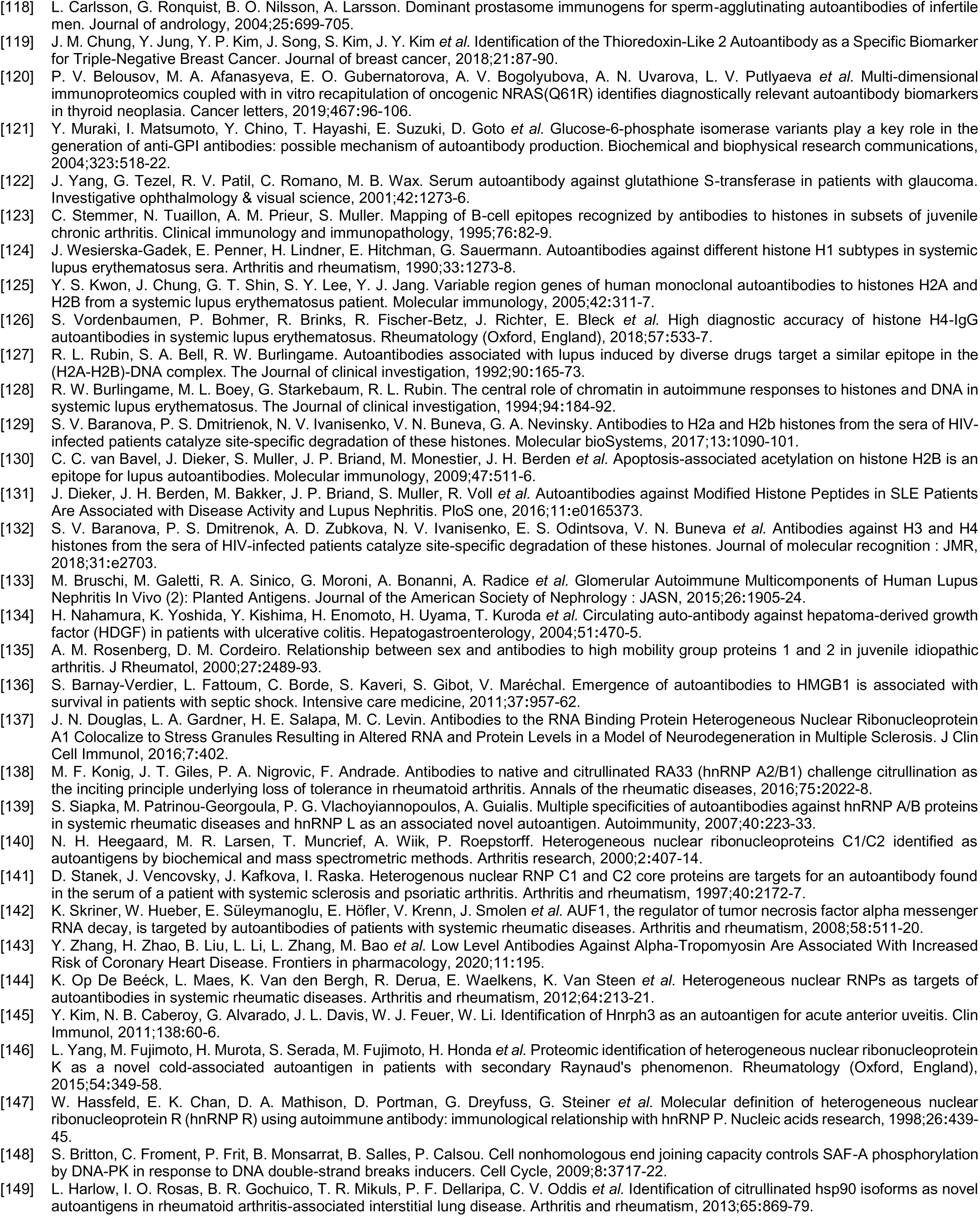
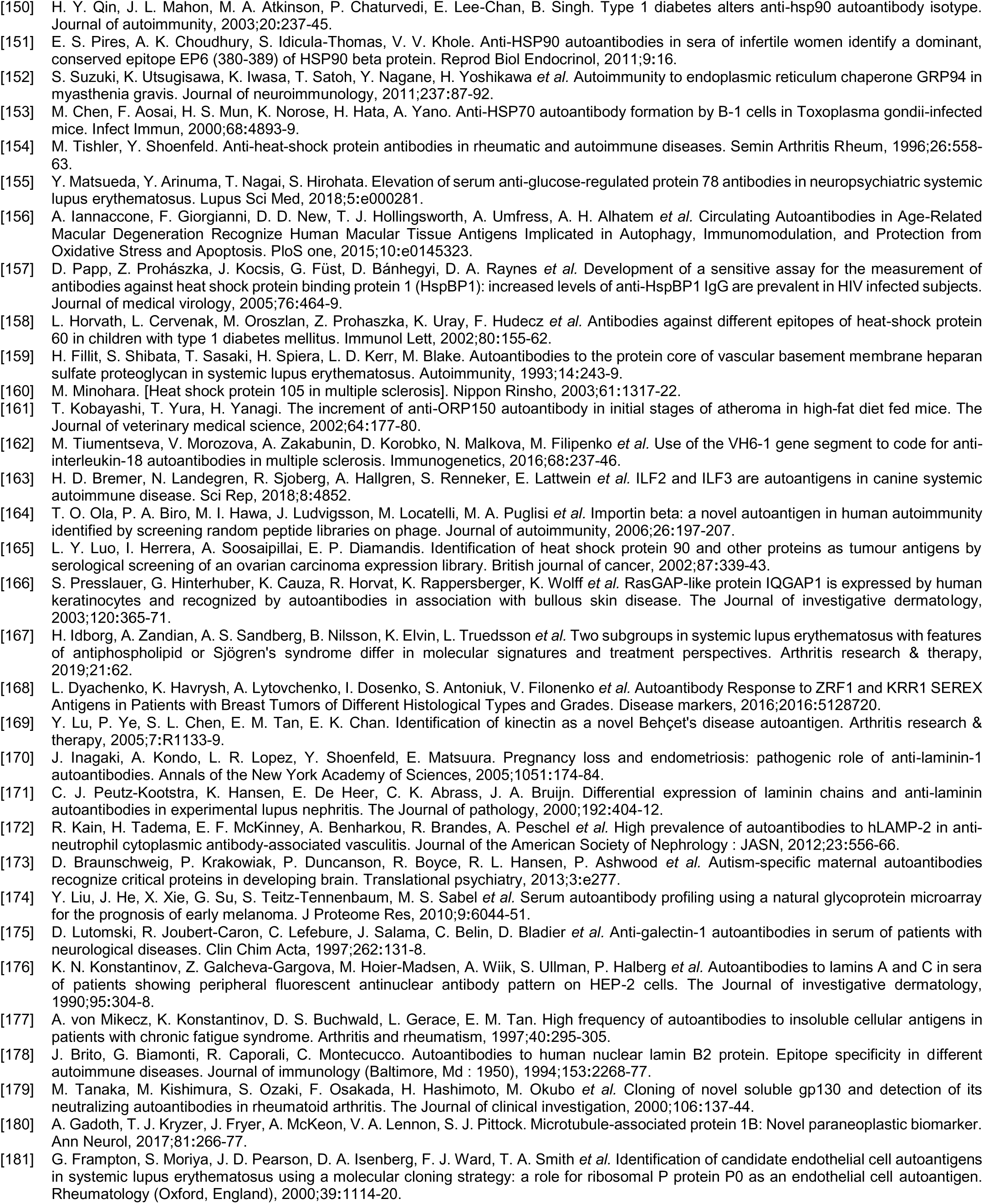
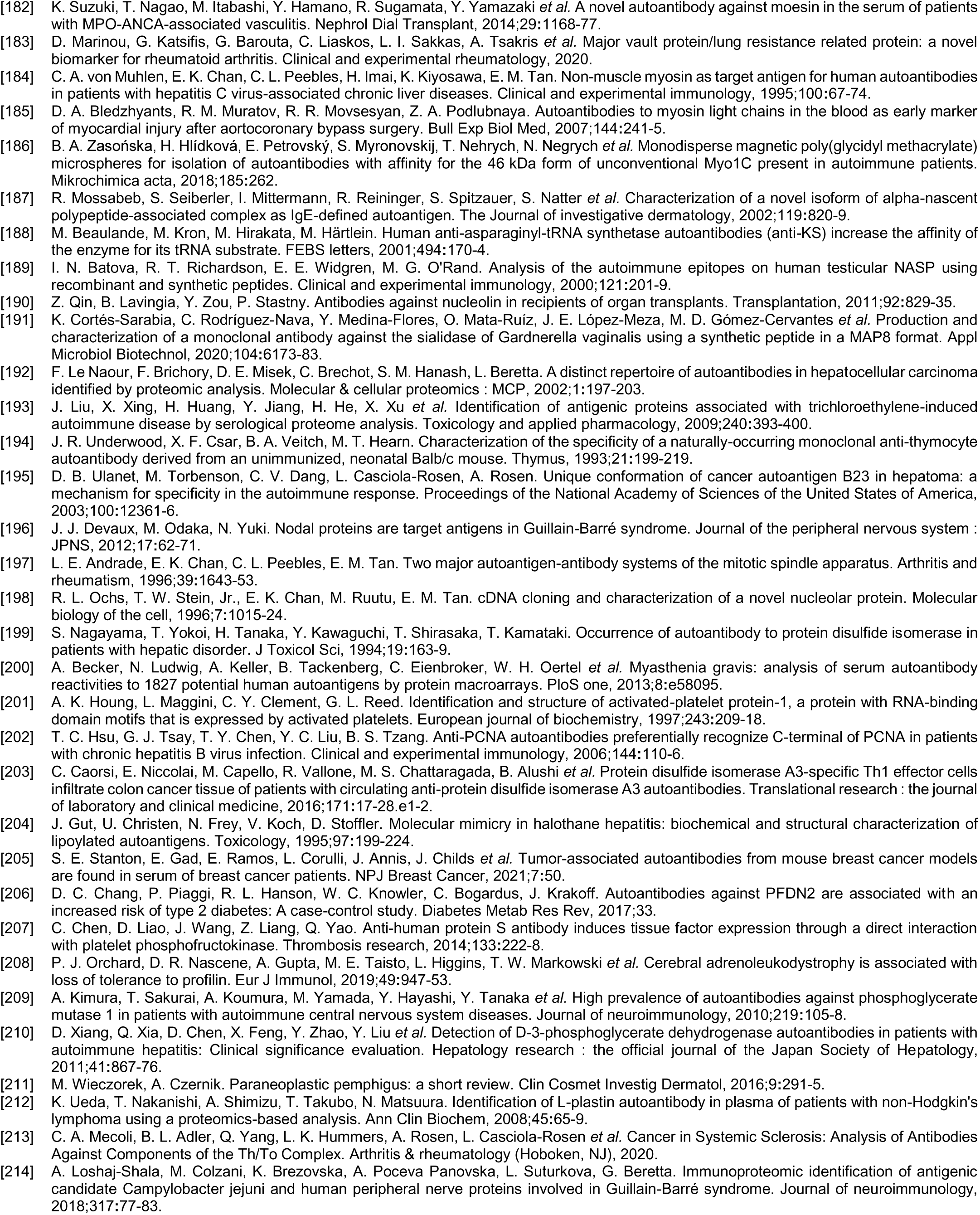
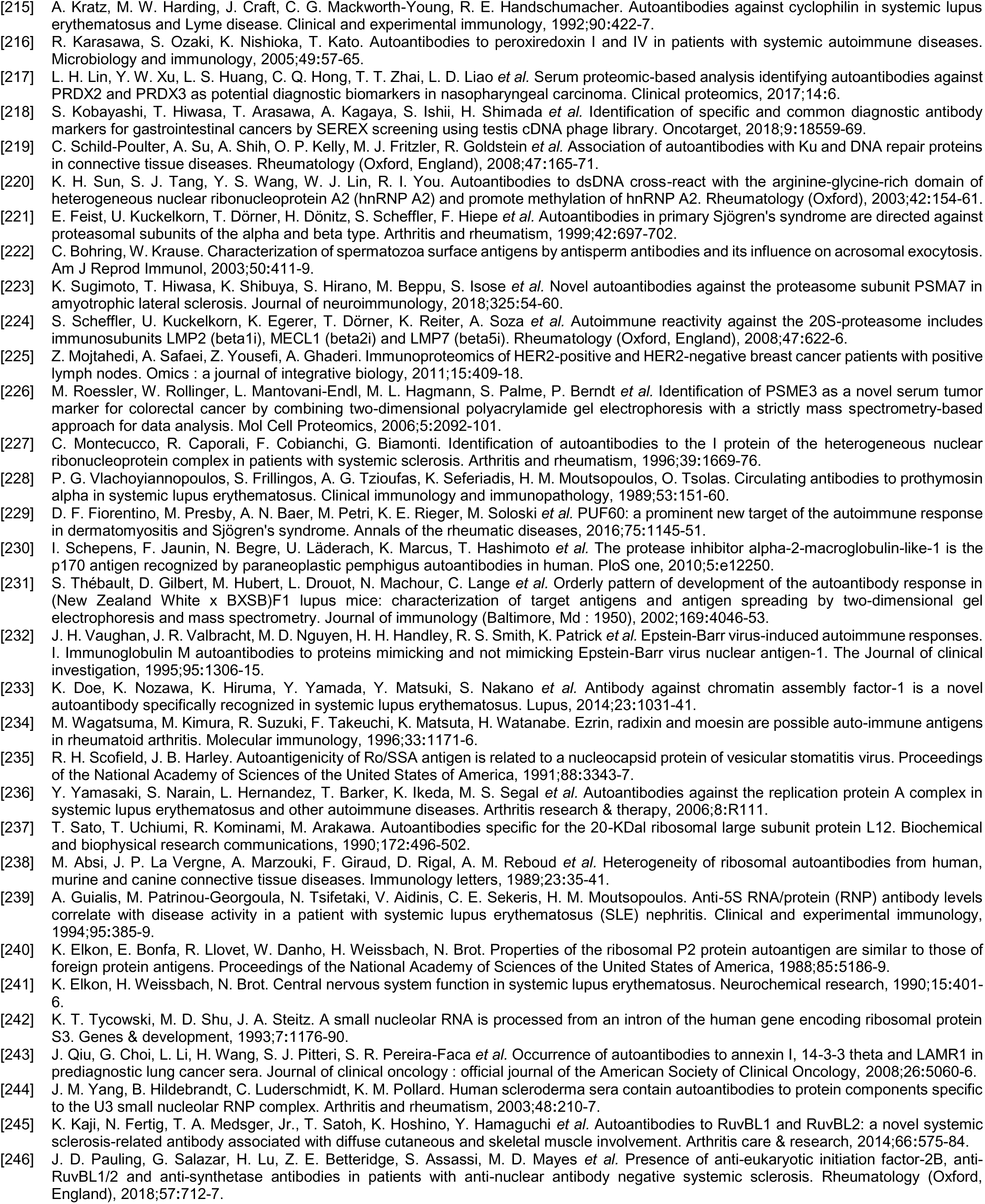
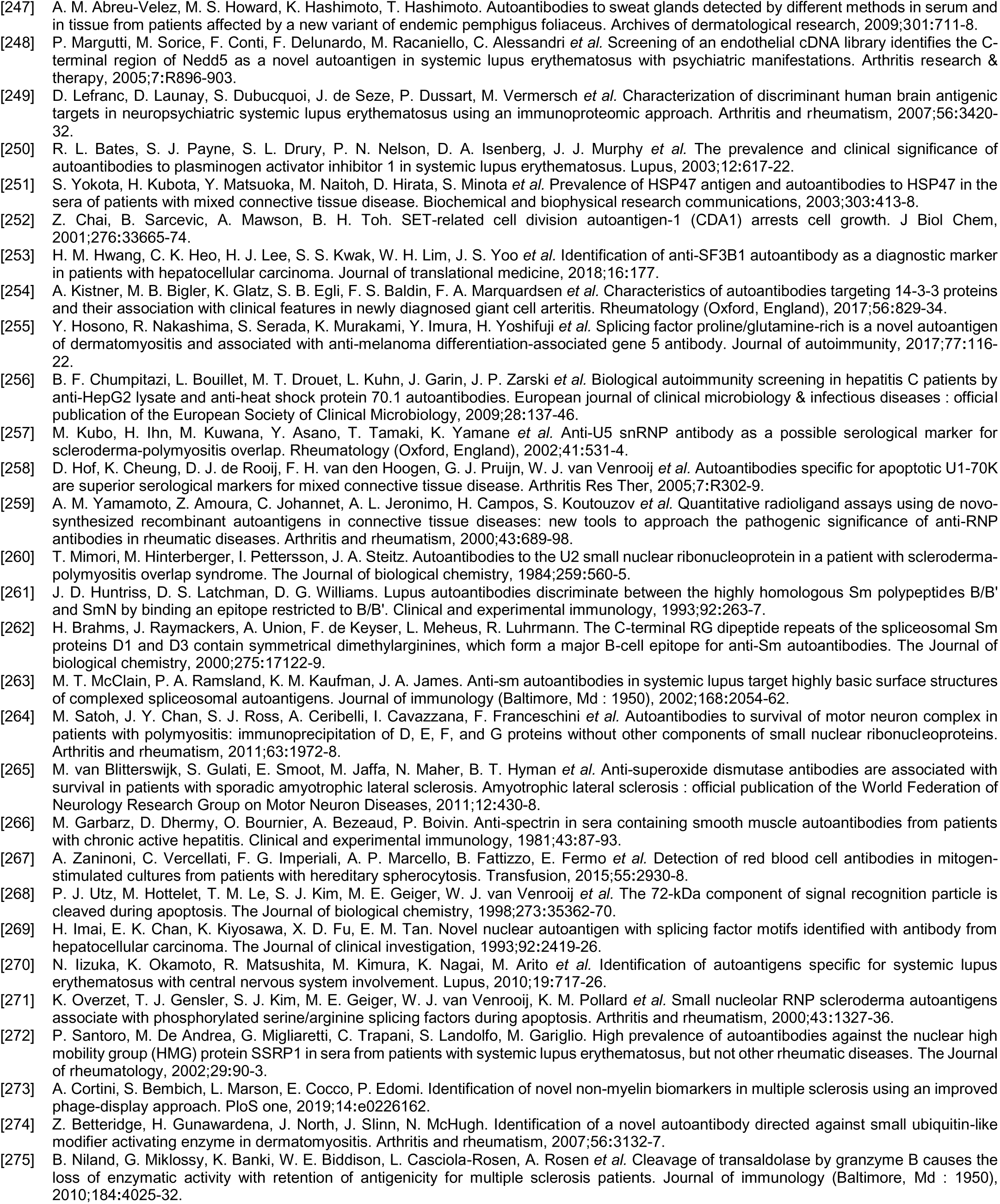
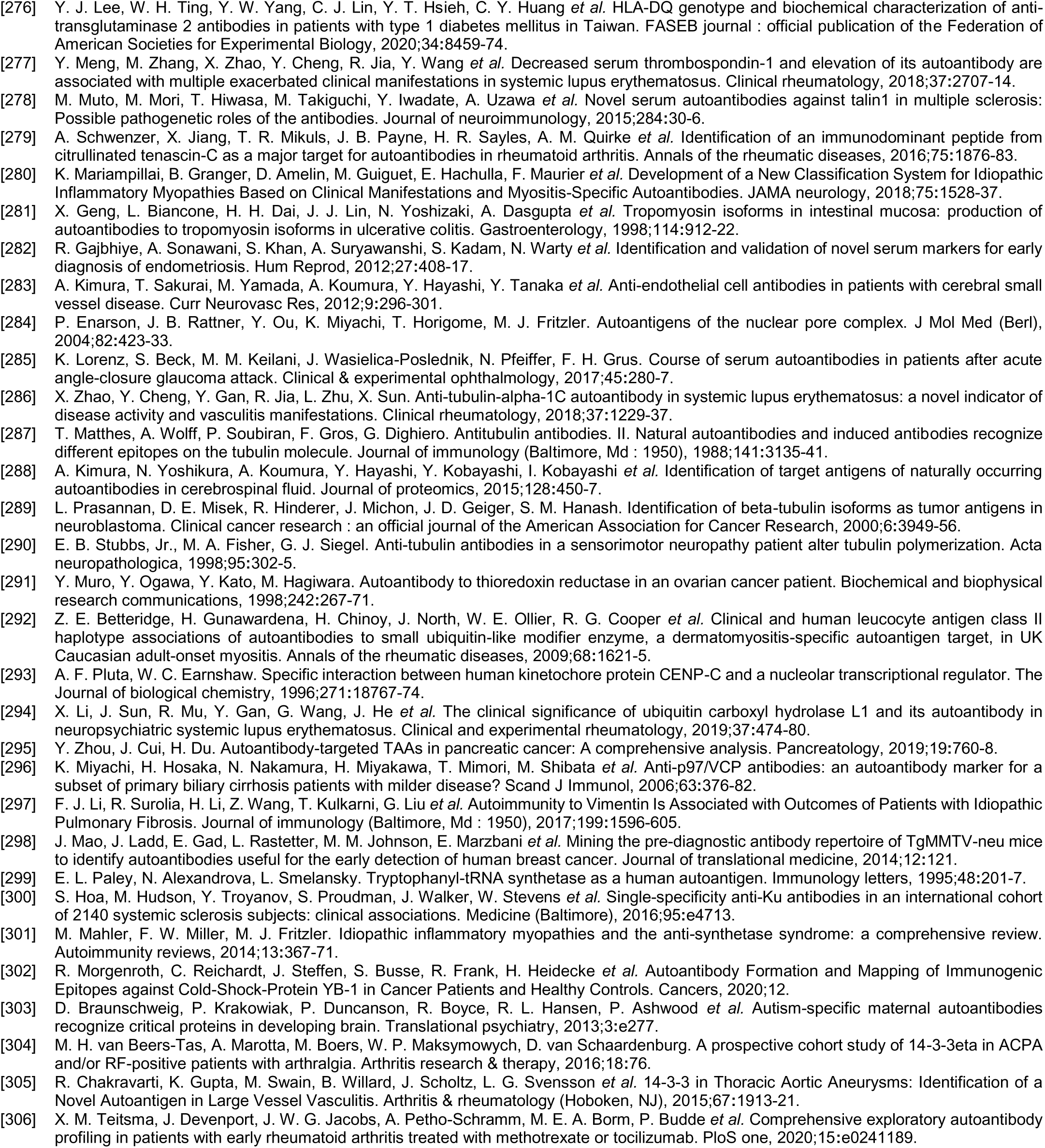
Autoantigens identified by DS-affinity and their alterations in SARS-CoV-2 infection

The master autoantigen-ome contains clusters of protein families, including 56 ribosomal proteins, 27 proteasome subunits, 19 heterogeneous ribonucleoproteins, 17 splicing factors, 17 ATP-dependent RNA helicase subunits, 16 eukaryotic translation initiation factors, 16 histones, 16 aminoacyl-tRNA synthases, 12 heat shock proteins, 9 elongation factors, 9 small nuclear ribonucleoproteins, 8 T-complex protein 1 subunits, and 7 14-3-3 proteins. In addition, there are multiple isoforms of numerous proteins, such as actin, tropomyosin, myosin, collagen, tubulin, and annexin.

The 751 confirmed and putative autoAgs are highly connected and have significantly more interactions than what would be expected for a random set of proteins of similar size drawn from the genome (exhibiting 6,936 interactions vs. 3,596 expected with the highest confidence level cutoff; enrichment p value <1.0e-16) as per protein-protein interaction analysis in STRING [14] (Fig. 2). The 400 confirmed autoAgs also form a similar, strong interacting network (exhibiting 2,758 interactions vs. 1,269 expected; enrichment p value <10e-16) (Fig. 3). The tight connections within the autoAg network suggest that these proteins are biologically connected, and given that they are all identified by DS-affinity, the autoAg protein networks offer a glimpse of the biological roles and functions of DS that await further investigation.

**Fig. 2.**
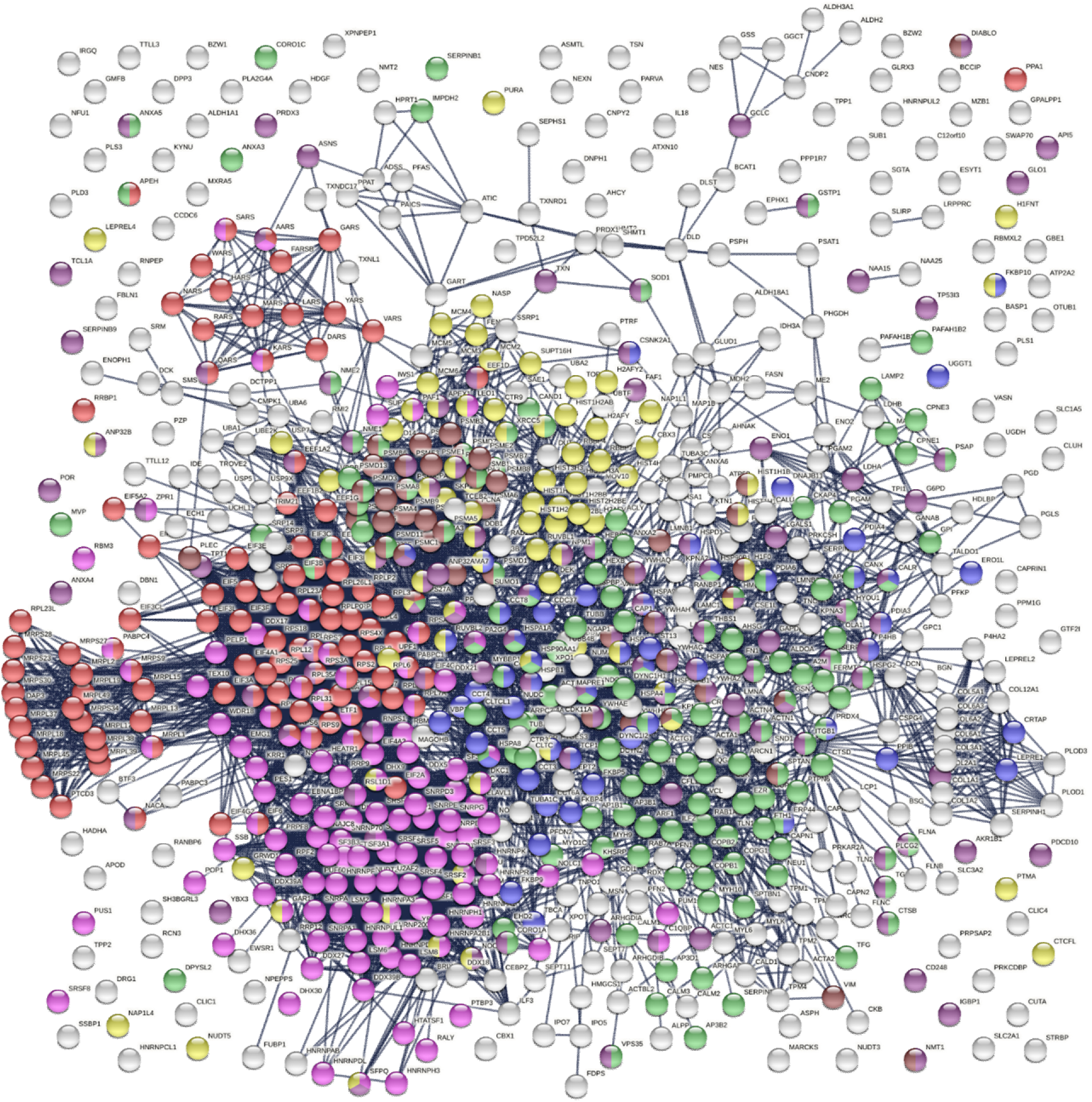
The master autoAg-ome of 751 DS-affinity proteins identified from 6 cell types forms a highly interacting connected network. Lines represent protein-protein interactions with the highest confidence cutoff. Colored proteins are associated with translation (104 proteins, red), RNA processing (120 proteins, pink), protein folding (53 proteins, blue), vesicle-mediated transport (141 proteins, green), chromosome organization (76 proteins, yellow), regulation of cell death (110 proteins, dark purple), and apoptosis (46 proteins, brown).

**Fig. 3.**
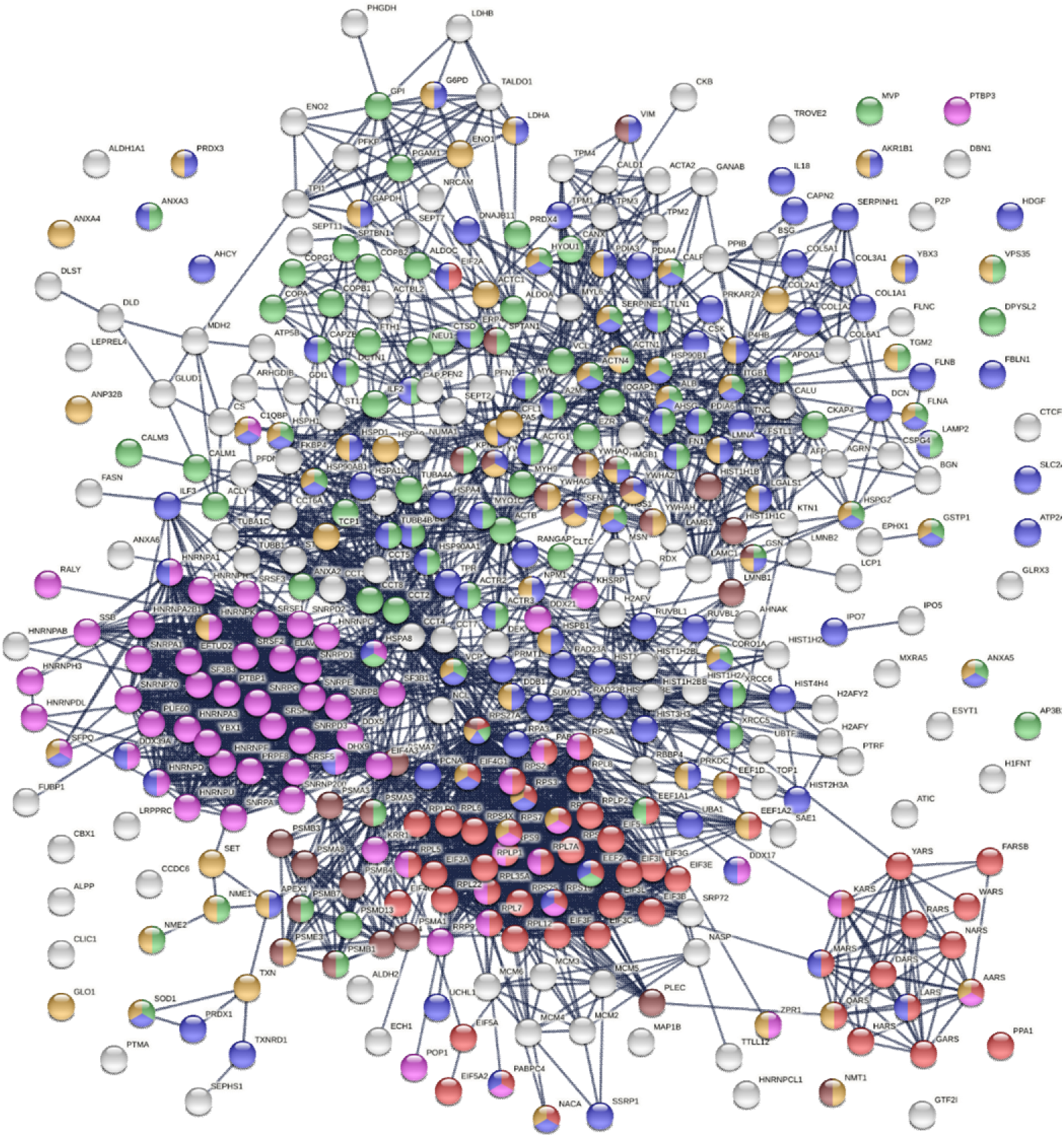
Protein interaction network of the 400 confirmed autoAgs. Lines represent protein-protein interactions with highest confidence. Colored proteins are associated with translation (57 proteins, red), RNA processing (65 proteins, pink), vesicle-mediated transport (89 proteins, green), response to stress (125 proteins, blue), regulation of cell death (74 proteins, amber), and apoptosis (28 proteins, brown).

The 751-protein master autoantigen-ome is significantly associated with many biological processes and pathways, most notably translation, RNA processing, RNA splicing, protein folding, vesicle-mediated transport, chromosome organization, regulation of cell death, and apoptosis (Figs. 2 and 4). The 400 confirmed autoAgs are similarly significantly associated with the same processes and pathways (Fig. 3). In addition, these proteins are associated with numerous other processes, e.g., mRNA metabolic process, peptide metabolic process, establishment of localization in the cell, intracellular transport, interspecies interaction between organisms, viral process (infection and virulence), symbiotic process, and response to stress (Figs. 2-4). Hierarchical clustering [15] of the top 50 enriched Gene Ontology Biological Processes reveals RNA processing, particularly RNA splicing, to be the most noticeable (Fig. 4).

**Fig. 4.**
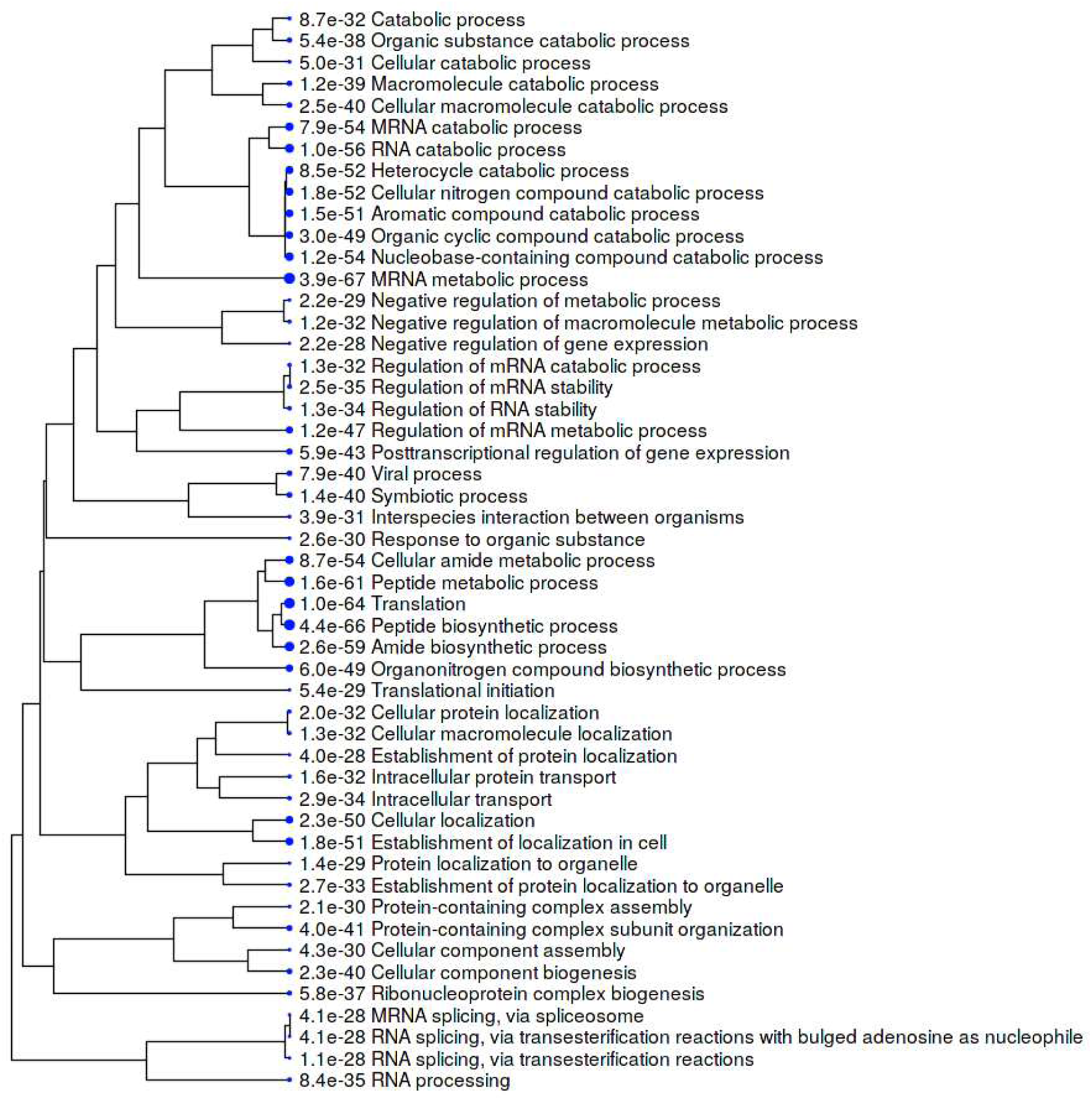
Hierarchical clustering of the top 50 GO Biological Processes associated with the master autoantigen-ome of 751 DS-affinity autoAgs. Bigger blue dots indicate more significant p values.

### The COVID-19 autoantigen-ome

To find out how many autoAgs in the autoantigen-ome are potentially affected by SARS-CoV-2 infection, we looked for them in currently available multi-omic COVID data compiled by Coronascape [16–37]. Remarkably, 657 (87.5%) proteins of the 751-member master autoantigen-ome are found to be affected in SARS-CoV-2 infection (Table 1 and Supplemental Table 1). Among them, 109 proteins were found up-regulated only, 176 were found down-regulated only, and 343 were found both up- and down-regulated at protein and/or RNA levels in virally infected cells or COVID-19 patients (Table 1 and Fig. 6). In addition, 191 potential autoAgs were found in the interactomes of different SARS-CoV-2 viral component proteins, meaning that they may directly or indirectly interact with the virus.

The 657-member COVID autoantigen-ome is also a highly interacting protein network (Fig. 5). Not surprisingly, these proteins are significantly associated with processes that are crucial in viral infection, e.g., RNA processing, mRNA metabolic process, regulation of mRNA stability, translation, peptide biosynthetic process, protein folding, intracellular transport, vesicle-mediated transport, regulated exocytosis, symbiont process, and interspecies interaction between organisms, response to stress, regulation of cell death, and apoptosis (Fig. 5). We also analyzed the 109 up-only and the 176 down-only protein networks separately. Both networks are significantly associated with translation, RNA processing and splicing, and the proteasome, which further illustrates that these processes are perturbed by the viral infection (Fig. 6).

**Fig. 5.**
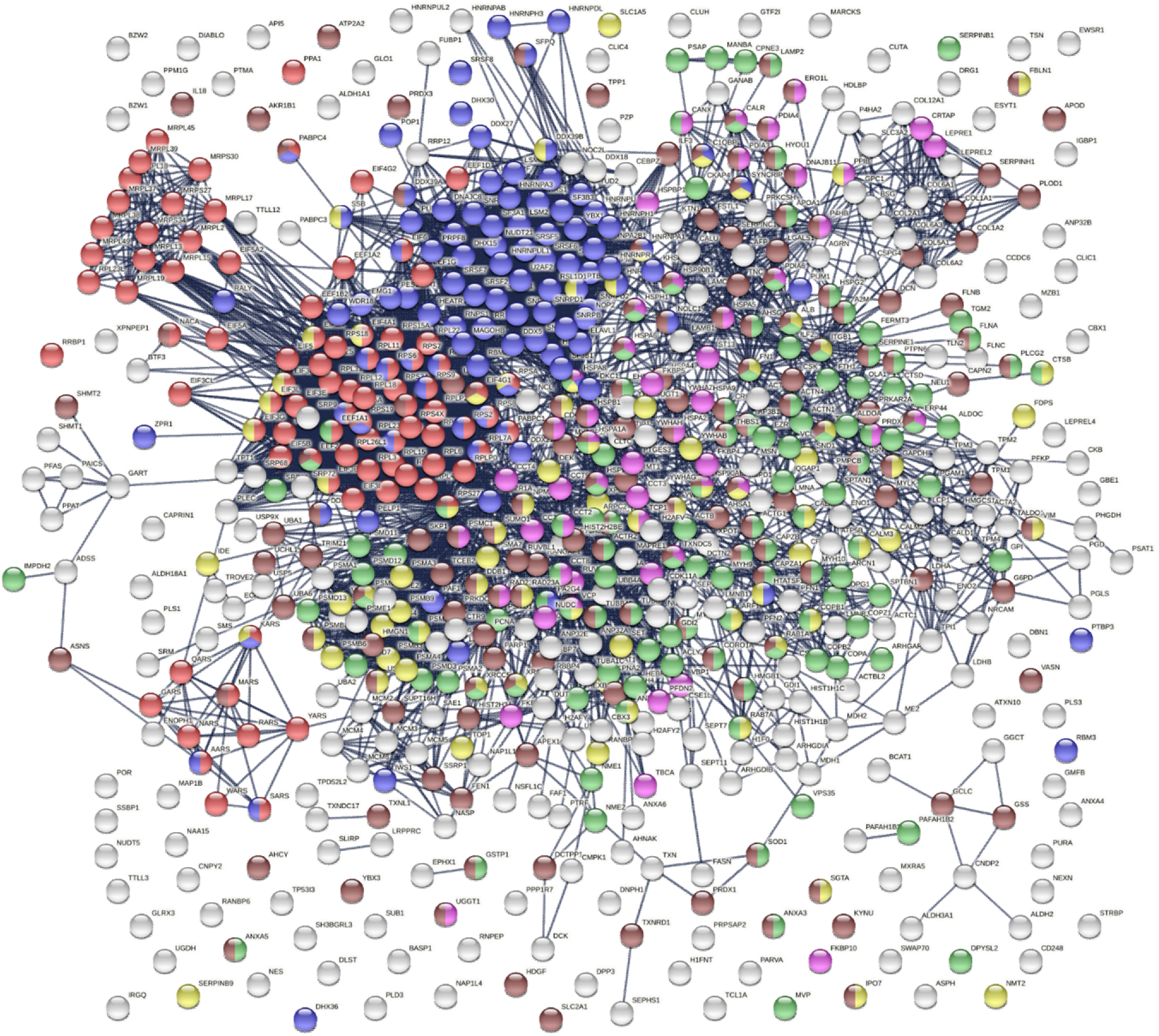
The COVID autoantigen-ome of 657 autoAg candidates. Lines represent protein-protein interactions with highest confident level. Colored proteins are associated with translation (87 proteins, red), RNA processing (103 proteins, blue), protein folding (51 proteins, pink), symbiont process (78 proteins, yellow), vesicle-mediated transport (125 proteins, green), and response to stress (161 proteins, brown).

**Fig. 6.**
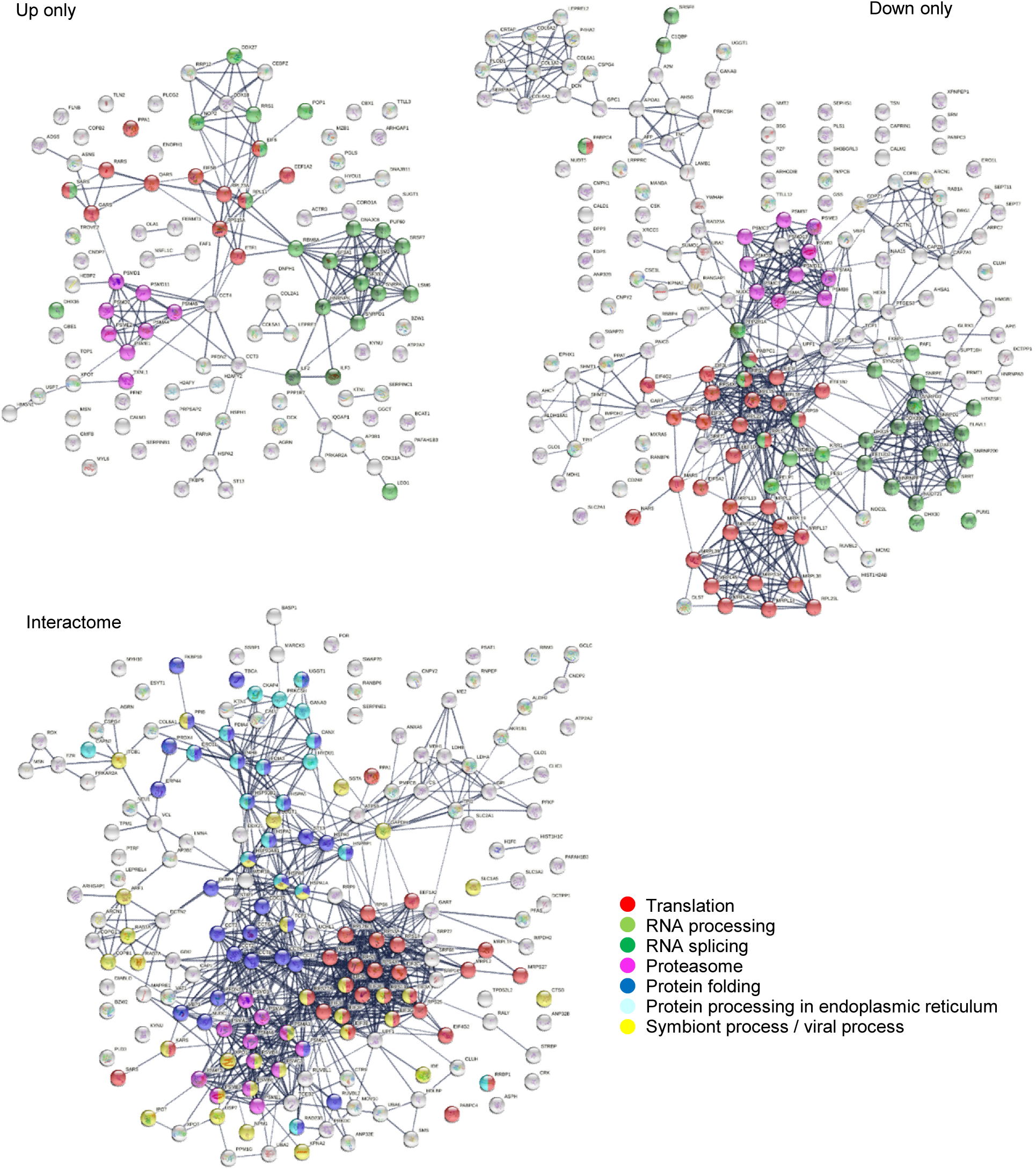
COVID-affected autoAgs that are found up-regulated only, down-regulated only, or interacting with SARS-Cov-2 proteins. Note the significant enrichment of proteins associated with translation, RNA processing and splicing, and other processes.

Translation is an essential step in viral replication and mRNA vaccine action. DS-affinity identified 19 eukaryotic translation initiation factors, with 15 thus far being confirmed autoAgs (Table 1). In particular, 8 of the 13 subunits of the human eIF3 complex were found in the interactome of the NSP1 protein of SARS-CoV-2, and all 8 are known autoAgs (Table 1). eIF3 is essential for the most forms of cap-dependent and cap-independent translation initiation and stimulates nearly all steps of translation initiation, as well as other phases of translation such as recycling. eIF3 functions in a number of prominent human pathogens, e.g., HIV and HCV; and the present finding indicates that eIF3 also functions in SARS-CoV-2 infection.

Among the 657 COVID-affected DS-affinity proteins, 369 (56%) are thus far confirmed autoAgs, accounting for 92% of the 400 confirmed autoAgs of the master autoantigen-ome. This vast number of perturbed autoAgs demonstrates that COVID-19 could lead to a wide variety of autoimmune diseases. For example, 42 autoAgs are associated with the myelin sheath and many are associated with other components of the nervous system, as we have described previously, which may help explain a myriad of neurological symptoms caused by COVID-19 [1]. As another example, 11 autoAgs are related to stress fibers (contractile actin filament bundles consisting of short actin filaments with alternating polarity) and 25 proteins are associated with myofibrils (contractile elements of skeletal and cardiac muscle), which may explain various muscular and cardiomuscular sequelae of COVID-19.

A few autoAgs also interact with multiple viral proteins of SARS-CoV-2, suggesting that they play important roles in COVID-19 and merit further investigation. For example, ESYT1 and MOV10 interact with 12 viral proteins, CALU interacts with 11, HSPA5 interacts with 9, COPG1 and ARHGAP1 interact with 8, PLD3 and MARCKS interact with 7, and IDE interacts with 6 viral proteins (Table 1). PLD3 (a phospholipase) influences the processing of amyloid-beta precursor protein and is associated with spinocerebellar ataxia and Alzheimer’s disease. IDE (insulin-degrading enzyme) degrades intracellular insulin and is associated with diabetes.

### AutoAg coding gene characteristics and alternative splicing

To further understand the autoantigen-ome, we mapped the coding genes for 751 proteins of the master autoantigen-ome, and they are distributed over all chromosomes (Fig. 7). Since these include both confirmed and putative autoAgs, one may argue that some of the putative autoAgs may not be true and the gene characteristics may not be meaningful. Therefore, we also mapped the genes for the 400 confirmed autoAgs, and they are similarly distributed over all chromosomes (Fig. 7). For both confirmed and putative autoAgs, coding gene prevalence is significantly higher on chromosomes 11, 12, 17, and 19, lower on chromosome 18, and almost absent on chromosome Y (Fig. 7). Various cluster loci are noticeable, e.g., on chromosomes 1, 11, 12, 17, and 19.

**Fig. 7.**
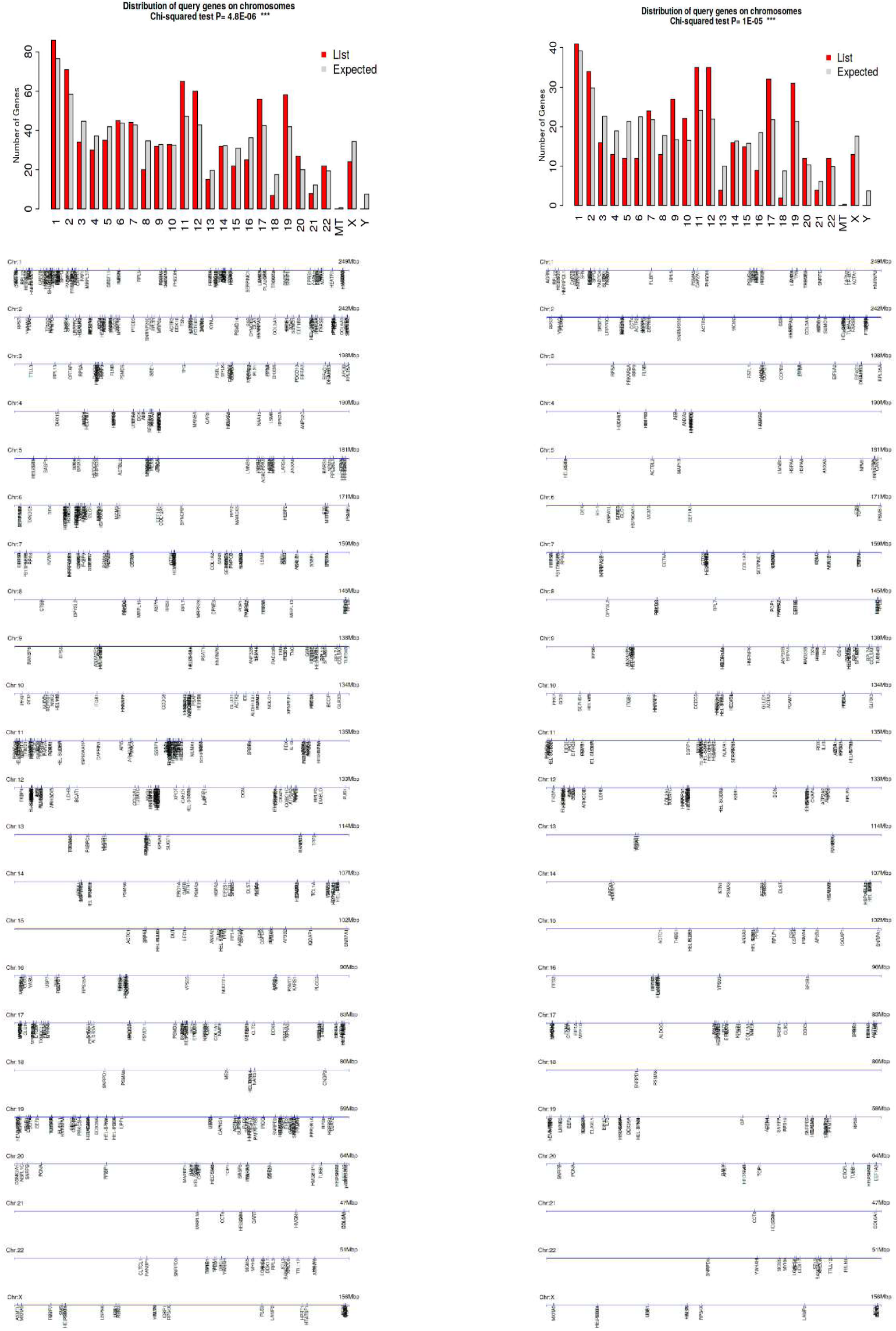
Distribution of autoAg coding genes by chromosomes. Left: 751 confirmed and putative autoAgs. Right: 400 confirmed autoAgs only.

Intriguingly, autoAg coding genes contain significantly larger numbers of exons than expected, with the majority containing at least 4 exons (Fig. 8). The number of transcript isoforms per coding gene is also significantly skewed towards higher numbers, and those with ≥6 isoforms are particularly dominant. Furthermore, the lengths of coding sequence, transcript, and 3’ and 5’-UTR of autoAg coding genes are skewed towards shorter sizes relative to the distribution of all coding genes (Fig. 8). We also examined the coding genes of the 400 confirmed autoAgs, and they show similar dominance in higher number ofexons and -isoforms, shorter transcripts, and shorter 3’-UTR lengths (Fig. 8).

**Fig. 8.**
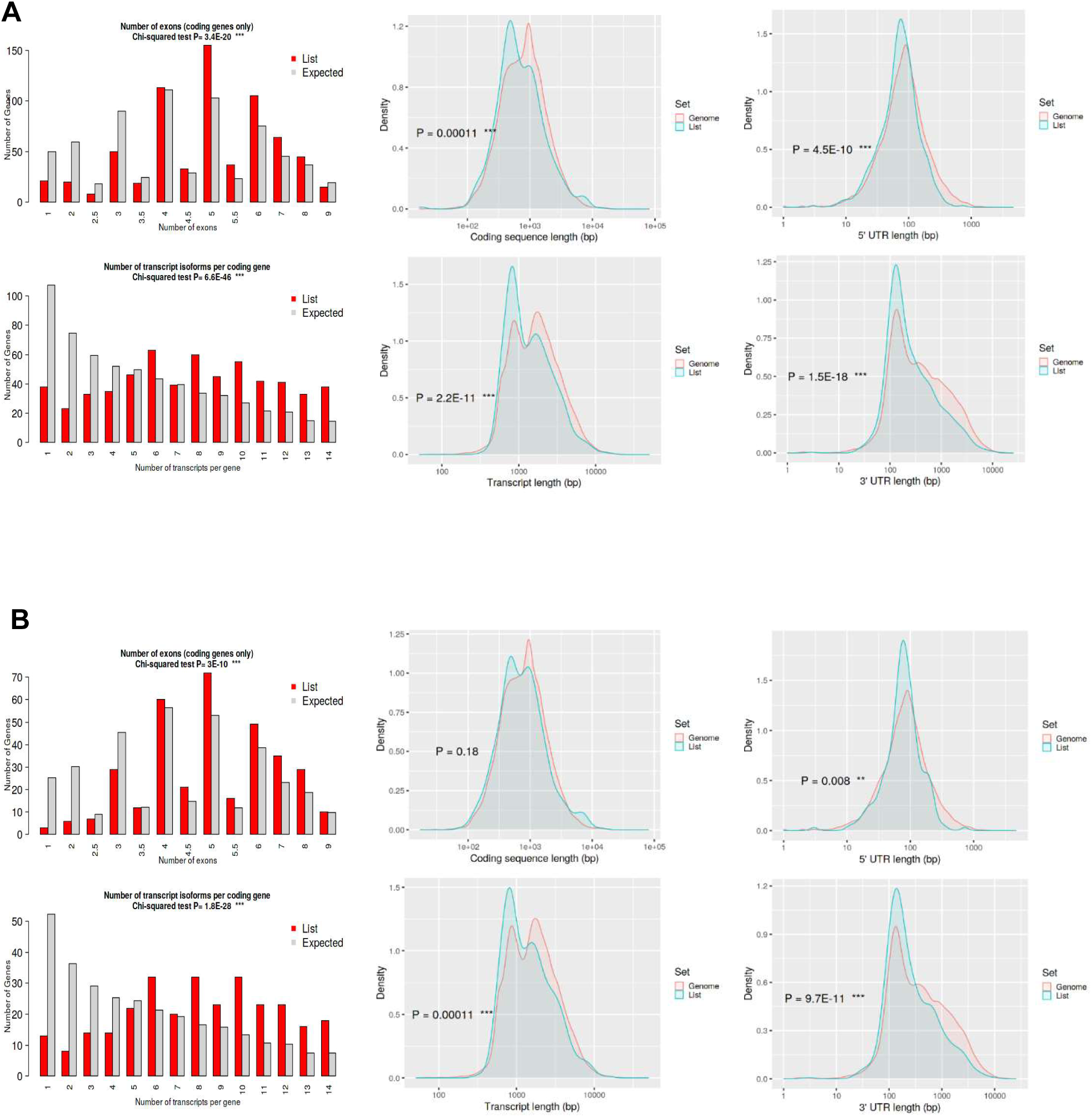
Characteristics of the autoAg coding genes compared with the rest in the genome. Differences are evaluated with Chi-squred and Student’s t-tests. (A) 751 confirmed and putative autoAgs. (B) 400 confirmed autoAgs.

The predominance of multiple exons and transcript variants suggests a role for RNA processing and alternative splicing in the origination of autoAgs. For genes with multiple exons, alternative splicing can yield a range of unique protein isoforms by varying the exon composition. Curiously, numerous components of the splicing machinery are well-known nuclear autoAgs. In fact, this study identified 120 potential autoAgs associated with RNA processing and 70 potential autoAgs associated with RNA splicing (Table 1 and Figs. 2-3). The majority of these have been found to be affected by SARS-CoV-2 infection (Figs. 5-6).

During splicing, a group of snRNPs (small nuclear ribonucleoproteins) bind to the intron of a newly formed pre-mRNA and splice it to result in a mature mRNA. Ten snRNP autoAgs are identified by DS-affinity, 8 of which have been found to be affected by SARS-CoV-2 infection (Table 1). During splicing, snRNAs undergo conformational rearrangements that are catalyzed by the DEAH/DEAD box superfamily of RNA helicases. 11 such helicases are identified by DS-affinity, and 10 have been found to be affected by the viral infection (Table 1). Serine/arginine-rich splicing factors, such as SRSF1 (also known as alternative splicing factor 1), are sequence-specific splicing factors involved in pre-mRNA splicing. 9 SRSF proteins are identified by DS-affinity, with 7 found to be affected by the viral infection. Seven additional splicing factors are identified by DS-affinity (e.g., poly(U)-binding splicing factor PUF60), with all found to be affected by SARS-CoV-2 infection. Heterogeneous nuclear ribonucleoproteins (hnRNPs) play various roles in gene transcription and post-transcriptional modification of pre-mRNA, e.g., binding pre-mRNAs to render splice sites more or less accessible to the spliceosome and suppressing RNA splicing at a particular exon. 19 hnRNP proteins are identified by DS-affinity, with 17 found affected by SARS-CoV-2 infection.

The large number of autoAgs of the RNA splicing machinery and their involvement in SARS-CoV-2 infection provide support to the notion that viral infections exploit alternative splicing. It is logical to speculate that viruses hijack the splicing machinery to force the host to synthesize virus-beneficial protein isoforms and thereby reprogram the host cellular protein network so that the virus can survive and replicate. It is also plausible that protein isoforms from virus-induced alternative splicing are recognizable by our immune system as unusual and non-self and hence may trigger an (auto)immune response.

Various studies have reported alternative splicing among autoAgs. For example, an informatics analysis of 45 autoAgs showed that alternative splicing occurred in 100% of the transcripts, which was significantly higher than the ∼42% rate observed in a randomly selected set of 9,554 gene transcripts. Furthermore, 80% of the transcripts underwent non-canonical alternative splicing, which was significantly higher than the <1% rate in randomly selected human gene transcripts [38]. As another example, Ro52/SSA is one of the autoAg targets strongly associated with the autoimmune responses in mothers whose children have manifestations of neonatal lupus. The gene for full-length Ro52 spans 10 kb of DNA and contains 7 exons, and an alternatively spliced transcript encoding a novel autoAg expressed in the fetal and adult heart has been identified [39]. In a patient with primary Sjörgren syndrome, an alternative mRNA variant of the nuclear autoAg La/SSB was found to result from a promoter switch and alternative splicing [40].

### Common autoAgs associated with cell stress and apoptosis

We have consistently found that DS binds apoptotic cells regardless of cell type [6, 8]. To figure out which molecules are involved in this affinity, we searched for DS-affinity proteins shared in all 6 human cell lines of this study and found 39 autoAg candidates (Fig. 9). These include 9 ER chaperone complex proteins, 5 14-3-3 proteins, 3 hnRNPs, and 3 tropomyosin proteins. All are known autoAgs except for ANP32A and YWHAB (14-3-3 alpha/beta). Given that ANP32A’s paralog ANP32B and 5 other 14-3-3 isoforms are known autoAgs, it is likely they are also true autoAgs. Remarkably, several classical ANA (antinuclear antibody) autoAgs that define systemic autoimmune diseases are among the autoAgs found in the DS-affinity proteomes of all 6 human cell lines, including histone H1 and H4, SSB (lupus La), XRCC5/Ku80, XRCC6/Ku70, and PCNA. Because these autoAgs are commonly found in apoptotic cells, it is not surprising that autoimmune responses targeting these autoAgs tend to be systemic; in other words, they all are potential markers of systemic autoimmune diseases.

**Fig. 9.**
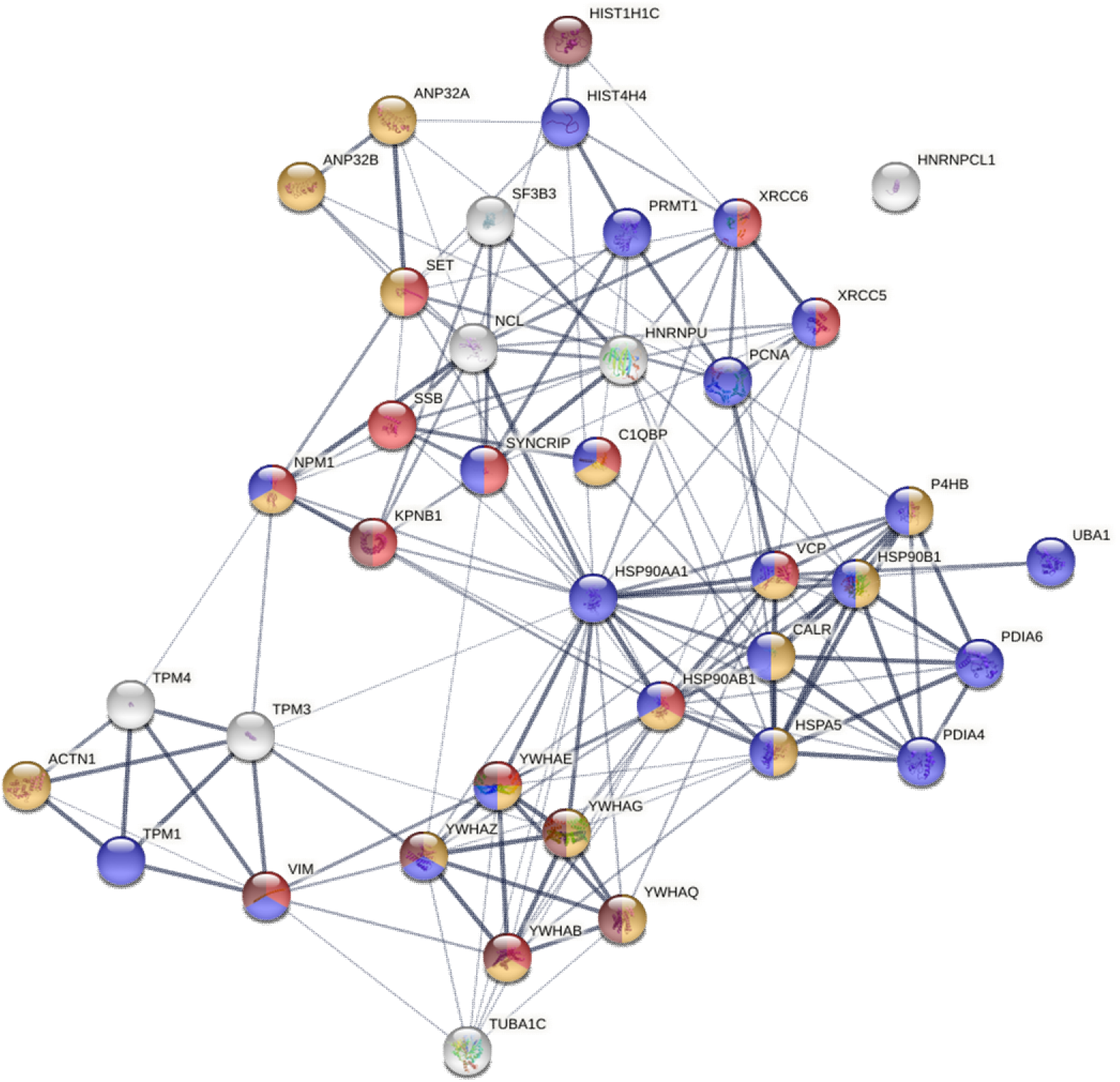
Common autoAgs identified from all six cell types examined in this study. Colored are proteins associated with viral infection (13 proteins, red), regulation of apoptotic process (17 proteins, amber), response to stress (22 proteins, blue), and apoptosis (8 proteins, brown).

Based on GO Biological Process and Reactome Pathway analysis, 22 of the common autoAgs are associated with cellular responses to stress, 17 are associated with regulation of apoptotic processes, and 8 are markers of apoptosis (Fig. 9). Moreover, these common autoAgs are involved in chromosome organization (ANP32A, ANP32B, H1-2, H4, KPNB1, NPM1, PCNA, SET, XRCC5, XRCC6), cytoskeleton organization (ACTN1, CALR, TPM1, TPM3, TPM4, TUBA1C, VIM), and mitochondrial membrane organization (YWHAB, YWHAE, YWHAG, YWHAQ, YWHAZ). These findings reveal that apoptosis is accompanied by reorganization of the nucleus, mitochondria, and cytoskeleton.

Furthermore, 37 of the 39 common autoAgs were altered in SARS-CoV-2 infection. Based on GO Biological Process analysis, 13 of these proteins are involved in viral processing, namely, KPNB1, C1QBP, HSP90AB1, NPM1, SYNCRIP, SET, SSB, XRCC5, XRCC6, VCP, VIM, YWHAB, and YWHAE. These findings further support our model of linking viral infection to autoimmunity, with viral infections leading to host cell stress, cell death, autoimmune reactions, and eventually autoimmune diseases (Fig. 1).

### UBA1, X-inactivation escape, and female predilection of autoimmunity

Among the above common autoAgs, UBA1 (or UBE1, ubiquitin-like modifier-activating enzyme 1) plays an essential role in dead cell clearance. UBA1 catalyzes the first step in ubiquitination – the “kiss of death” – that marks cellular proteins for degradation. It has long been speculated that dysregulation of apoptotic pathways and dysfunctional clearance of dead cells are among the main causes of autoimmunity, which is in line with our findings [6, 8]. Apoptosis also directly contributes to the maintenance of lymphocyte homeostasis and the deletion of autoreactive cells. Therefore, dysfunction of UBA1 could result in deficient clearance of apoptotic cells and aberrant autoimmunity.

Recently, UBA1 somatic mutations have been linked to a severe adult-onset autoinflammatory disease termed VEXAS syndrome [41]. A somatic mutation affecting methionine-41 in UBA1 results in a loss of the canonical cytoplasmic isoform of UBA1 and in the expression of a novel catalytically impaired isoform. Additionally, mutant peripheral blood cells show decreased ubiquitination and activated innate immune pathways.

Strikingly, UBA1 protein expression is found up-regulated at different time points of SARS-CoV-2 infection, whereas two deubiquitinating enzymes, USP9X and USP5, are down-regulated [33] (Supplemental Table 1). Furthermore, among the 657 proteins of the COVID autoantigen-ome, 178 have been found to be affected by ubiquitination (Fig. 10). They are most significantly associated with RNA metabolism and cellular response to stress. In addition, ubiquitination affects proteins involved in signaling by Rho GTPase, RNA splicing, translation, protein folding, nonsense-mediated decay, DNA damage stress-induced senescence, and the cytoskeleton. These findings underline the extensive involvement of ubiquitination in viral infection.

**Fig. 10.**
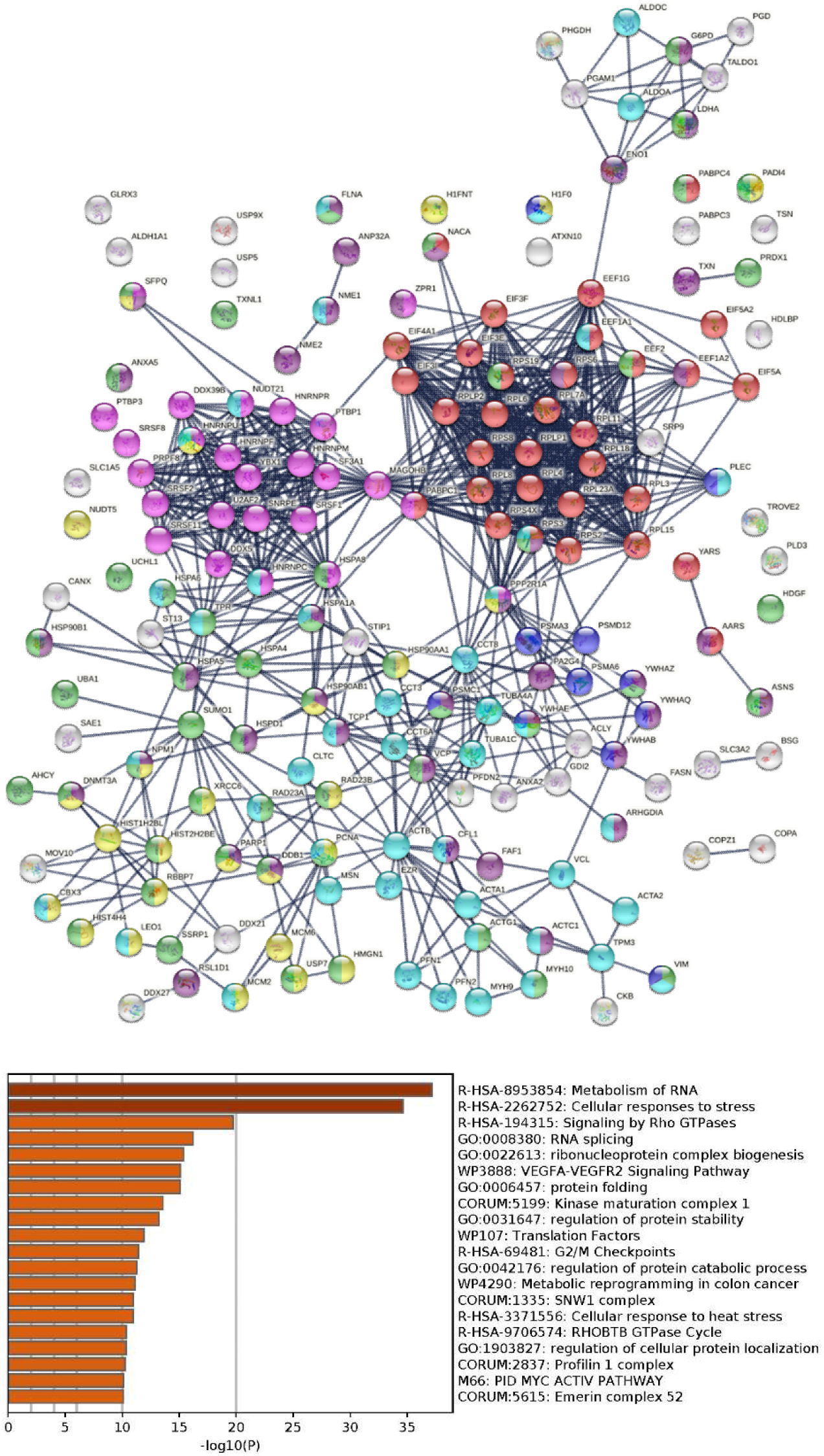
Top: Potential autoAgs affected by ubiquitination in SARS-Cov-2 infection (lines represent protein-protein interactions with the highest confidence). Colored are proteins associated with translation (32 proteins, red), RNA splicing (25 proteins, pink), regulation of cell death (40 proteins, dark purple), chromosome organization (26 proteins, yellow), response to stress (50 proteins, green), cytoskeleton (45 proteins, aqua), and apoptosis (11 proteins, blue). Bottom: Top 20 enriched processes and pathways associated with the ubiquitinated autoAgs .

UBA1 is coded by the *UBA1* gene located on the X chromosome with no homolog on the Y chromosome, and more importantly, *UBA1* can escape X-chromosome inactivation. *UBA1* appears to be protected against chromosome-wide transcriptional silencing by a chromatin boundary flanked by histone H3 modifications and CpG hypomethylation [42]. In human female fibroblasts, UBA1 mRNA is detected from both the active and inactive X chromosomes, and UBA1 is expressed in a large panel of somatic cell hybrids retaining inactive X chromosomes [43]. In human endothelial cells from dizygotic twins, UBA1 and a few other X-chromosome encoded proteins are expressed at higher levels in female cells [44]. UBA1 expression is estimated to be ∼ 60% from X-active alleles, 30% biallelic, and 10% from X-inactive alleles [45].

X-linked genes, particularly escape genes, contribute to sex differences. In women, about 15% of X-linked genes are bi-allelically expressed, and expression from the inactive X allele varies from a few percent to near equal to that of the active allele [46]. X-inactivation and escape may enhance phenotypic differences between females and males and may also enhance variability within females due to mosaicism from cells with the X-maternal or X-paternal inactivated and to a variable degree of escape from X-inactivation [46]. Aging, which is associated with telomere shortening, can relax X-inactivation and force global transcriptome alterations [47], which may lead to gene escape and altered expression of UBA1. Therefore, dysfunction of UBA1 due to X-inactivation escape may predispose women, particularly aging women, to increasing dysfunctional regulation of apoptosis and aberrant autoimmunity.

### Considerations for vaccine design based on Spike-protein via viral vectors or mRNAs

To understand the various rare but reported side effects from the currently available viral vector- and mRNA-encoded S-protein COVID vaccines, we searched for autoAgs that may interact with the spike protein of SARS-CoV-2 and found 15 autoAg candidates (Table 2). Of these, CALU, ESYT1, MOV10, and MARCKS may also interact with many other SARS-CoV-2 proteins as discussed earlier. Curiously, at least 2 of these are associated with blood clotting problems, and 5 are implicated in neurological disorders (Table 2). For example, CALU (calumenin) is a calcium-binding protein and is expressed in high levels in the heart, placenta, and skeletal muscle. CALU is associated with pharmacodynamics and response to elevated platelet cytosolic Ca^2+^, platelet degranulation, and Coumarin/Warfarin resistance. Warfarin is an anticoagulant (blood thinner) drug used to treat blood clots such as deep vein thrombosis and pulmonary embolism and to prevent stroke in people with heart problems such as atrial fibrillation, valvular heart disease or in people with artificial heart valves.

**Table 2.**
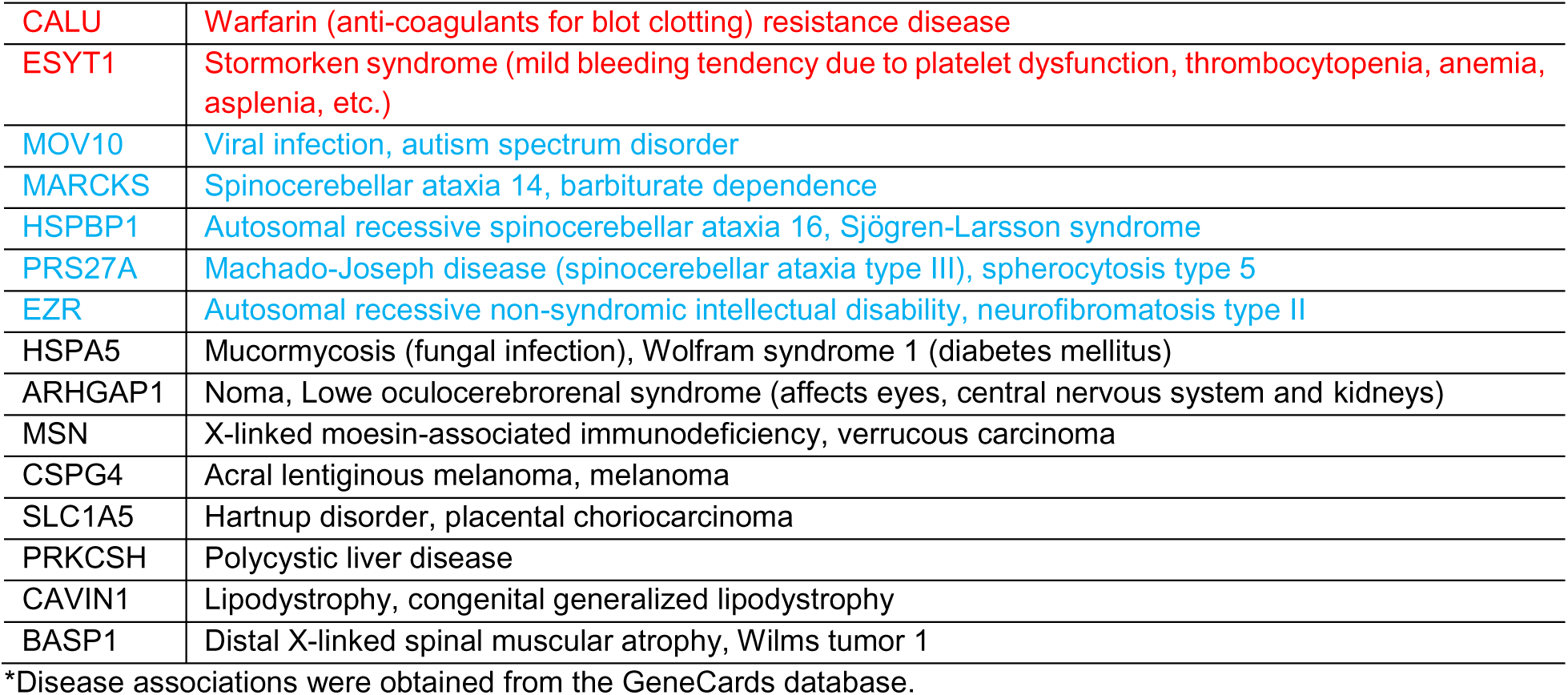
Diseases associated with potential SARS-CoV-2 spike protein-interacting autoAgs*

Although largely speculative at present, these potential S-protein-interacting autoAgs may provide partial explanations for the rare hematological, neurological, and muscular side effects reported for the currently available COVID vaccines (Table 2). Although it is known that S proteins are synthesized intracellularly following vaccination with mRNAs or viral vectors, many of the precise molecular steps remain unknown. In particular, how do these newly synthesized S proteins fold and are they glycosylated differently depending on the cell type that rakes up the mRNA or the viral vector? How does the newly synthesized S protein interact with other host cell components before being processed (or degraded) and presented to immune cells? For example, could the nascent S proteins interact with CALU or ESYT1 to cause blood clotting problems, could S protein interaction with HSPA5 contributes to fungal infection outbreaks as seen in India? These and many other questions await further investigation. This is of interest because mRNA and vector-based vaccines make use of a variety of cell types in vivo to produce the immunogen, whereas recombinant protein-based vaccines introduce the ex vivo prepared immunogen directly to the immune system.

In addition, this study identified a large number of autoAg candidates that are crucial for vector-based or mRNA vaccine action, including translation, RNA processing and metabolism, vesicles and vesicle-mediated transport, and protein processing and transport (Figs. 2-6). For example, the master autoantigen-ome contains 56 ribosomal proteins, 16 eukaryotic translation initiation factors, 16 aminoacyl-tRNA synthases/ligases, and 6 translation elongation factors, all of which are essential actors in translating mRNAs into proteins. There are also many autoAgs related to protein folding and post-translational protein modification, although it is not clear whether the S proteins are folded and post-translationally modified before being processed and presented to immune cells in the currently used mRNA or vector vaccines for COVID-19. These potential autoAgs may confer clues to understanding the observed rare adverse events and should help guide the future development of even safer vaccines.

## Conclusion

In this report, we compiled a master autoantigen-ome of 751 potential autoAgs, 657 of which are affected in SARS-CoV-2 infection, and 400 of which are confirmed autoAgs in a wide variety of autoimmune diseases and cancer. Our proposed model (Fig. 1) provides a plausible explanation for how a cascade of molecular changes associated with viral infection leads to cell stress, apoptosis, and subsequent autoimmune responses. The large number of autoAg candidates associated with SARS-CoV-2 infection provides a mechanistic rationale for the close monitoring of autoimmune diseases that may follow the COVID-19 pandemic. In addition, the coding gene characteristics of autoAgs described in this study provide further insights into the genetic origination of autoAgs. The significance of ubiquitination in apoptotic cell clearance and protein turnover and the X-linked escape expression of UBA1 might explain, in part, the predisposition of aging women to autoimmune diseases.

## Materials and Methods

### DS-affinity autoAg identification

Potential autoAgs were identified by DS-affinity from protein extracts from six human cell lines as previously described, including HFL1 fetal lung fibroblasts [1], A549 lung epithelial cells [2], HS-Sultan B-lymphoblasts [4], Wil2-NS B-lymphoblasts [7], Jurkat T-lymphoblasts [5], and HEp-2 fibroblasts [11].

### Autoantigen literature text mining

Each DS-affinity protein was verified as to whether it is a target of autoantibodies by an extensive literature search on PubMed. Search keywords included the MeSH keyword “autoantibodies”, the protein name or its gene symbol, or alternative names and symbols. Only proteins for which specific autoantibodies are reported in PubMed-listed journal articles were considered “confirmed” or “known” autoAgs in this study.

### COVID data comparison

DS-affinity proteins were compared with currently available COVID-19 multi-omic data compiled in the Coronascape database (as of 05/27/2021) [16–37]. These data have been obtained with proteomics, phosphoproteomics, interactome and ubiquitome studies, and RNA-seq techniques. Up- and/or down-regulated proteins or genes were identified by comparing cells infected vs. uninfected by SARS-CoV-2 or COVID-19 patients vs. healthy controls. Similarity searches were conducted to identify DS-affinity proteins that are similar to those found up- and/or down-regulated in the viral infection at any omic level.

### Protein network analysis

Protein-protein interactions were analyzed with STRING [14]. Interactions include both direct physical interaction and indirect functional associations, which are derived from genomic context predictions, high-throughput lab experiments, co-expression, automated text mining, and previous knowledge in databases. Each interaction is annotated with a confidence score between 0 (lowest) and 1 (highest), indicating the likelihood of an interaction to be true. Enrichment of pathways and processes were analyzed with Metascape [16], which utilize various ontological sources such as KEGG Pathway, GO Biological Process, Reactome Gene Sets, and Canonical Pathways. All genes in the genome were used as the enrichment background. Terms with a p value <0.01, a minimum count of 3, and an enrichment factor (ratio between the observed counts and the counts expected by chance) >1.5 were grouped into clusters based on their membership similarities. The most statistically significant term within a cluster was chosen to represent the cluster.

### Gene characteristic analysis

Gene characteristics were analyzed with ShinyGO [15]. ShinyGO is based on a large annotation database derived from Ensembl and STRING-db. The characteristics of the genes for the groups of autoAgs in this study were compared with the rest in the genome. Chi-squared and Student’s t-tests were run to see if the autoAg genes had special characteristics when compared with all other genes in the human genome.

## Authors’ contributions

JYW conducted the study and wrote the manuscript. MWR and VBR assisted with the study and manuscript preparation. MHR consulted on the study and edited the manuscript. All authors have approved the manuscript.

